# Exons as units of phenotypic impact for truncating mutations in autism

**DOI:** 10.1101/270850

**Authors:** Andrew H. Chiang, Jonathan Chang, Jiayao Wang, Dennis Vitkup

## Abstract

Autism spectrum disorders (ASD) are a group of related neurodevelopmental diseases displaying significant genetic and phenotypic heterogeneity^1–4^. Despite recent progress in understanding ASD genetics, the nature of phenotypic heterogeneity across probands remains unclear^5, 6^. Notably, likely gene-disrupting (LGD) *de novo* mutations affecting the same gene often result in substantially different ASD phenotypes. Nevertheless, we find that truncating mutations that affect the same exon frequently lead to strikingly similar intellectual phenotypes in unrelated ASD probands. Analogous patterns are observed for two independent proband cohorts and several other important ASD-associated phenotypes. We find that exons biased towards prenatal and postnatal expression preferentially contribute to ASD cases with lower and higher IQ phenotypes, respectively. These results suggest that exons, rather than genes, often represent a unit of effective phenotypic impact for truncating mutations in autism. The observed phenotypic effects are likely mediated by nonsense-mediated decay (NMD) of splicing isoforms, with autism phenotypes usually triggered by relatively mild (15-30%) decreases in overall gene dosage. We find that each gene with recurrent ASD mutations can be described by a parameter, phenotype dosage sensitivity (PDS), which characterizes the quantitative relationship between changes in a gene’s dosage and changes in a given disease phenotype. We further demonstrate analogous relationships between LGD mutations and changes in gene expression across human tissues. Therefore, similar phenotypic patterns may be also observed in multiple other systems and genetic disorders.

## Introduction

Recent advances in neuropsychiatric genetics^(7–10^and, specifically, in the study of autism spectrum disorders (ASD)^(11–14^have led to the identification of multiple genes and specific cellular processes that are affected in these diseases^11, 12, 14–16^. However, phenotypes usually associated with ASD vary considerably across autism probands^(1–4^, and the nature of this phenotypic heterogeneity is not well understood^5, 6^. Despite the complex genetic architecture of ASD^(17–22^, a subset of cases from simplex families, i.e. families with only a single affected child among siblings, are known to be strongly affected by *de novo* mutations with severe deleterious effects^14, 23, 24^. Interestingly, despite having relatively simpler genetic architecture, simplex autism cohorts often display as much phenotypic heterogeneity as more general ASD cohorts^(25–27^. This provides an opportunity for an in-depth exploration of the etiology of the autism phenotypic heterogeneity, at least for these cohorts, using accumulated phenotypic and genetic data. In this study we performed such an analysis, focusing on severely damaging, so-called likely gene-disrupting (LGD) mutations, which include nonsense, splice site, and frameshift variants. We explored genetic and phenotypic data collected in the Simons Simplex Collection (SSC)^28^and then validated our results using an independent ASD cohort from the Simons Variation in Individuals Project (VIP)^29^.

In this paper we investigated the effects of LGD mutations on cognitive and other important ASD-related phenotypes, including adaptive behavior, motor skills, communication, and coordination. These analyses allowed us to understand how the exon-intron structure of human genes contributes to observed phenotypic heterogeneity. We then explored the quantitative relationships between changes in gene dosage induced by nonsense-mediated decay (NMD) and the phenotypic effects of LGD mutations. To that end, we introduced a new genetic parameter quantifying how changes in a gene’s dosage affect specific autism phenotypes. Finally, we described how simple linear models of gene dosage can explain a substantial fraction of the phenotypic heterogeneity in the analyzed simplex ASD cohorts.

## Results

We first considered the impact of *de novo* LGD mutations on several well-studied cognitive phenotypes: full-scale (FSIQ), nonverbal (NVIQ), and verbal (VIQ) intelligence quotients^11, 14, 16^; these scores are normalized by age and standardized across a broad range of phenotypes^28^. We analyzed *de novo* mutations and the corresponding phenotypes of ASD probands for more than 2,500 families from SSC^28^. Notably, we found that the average IQ differences between probands with LGD mutations in the same gene were only slightly smaller than the IQ differences between all pairs of probands; the mean pairwise IQ difference for probands with mutations in the same gene was 25.7 NVIQ points, while the mean difference for all pairs of probands was 29.4 NVIQ points (∼12% difference, Mann-Whitney U one-tail test P = 0.14; Supplementary Table 1).

We next asked whether probands with LGD mutations at similar locations within the same gene resulted, on average, in more similar phenotypes (Supplementary Fig. 1). Indeed, IQ differences between probands with LGD mutations closer than 1000 base pairs apart were significantly smaller than the IQ differences between probands with more distant mutations; NVIQ average difference of 10.4 points for ≤ 1000 bp, NVIQ average difference of 28.6 points for > 1000 bp (MWU one-tail test *P* = 0.005). However, across the entire range of nucleotide distances between LGD mutations in the same genes, we did not observe either a significant correlation or a monotonic relationship between IQ differences and mutation proximity (NVIQ Spearman’s *ρ* = 0.1, *P* = 0.4; Mann-Kendall one-tail trend test *P* = 0.5).

To explain the observed patterns of phenotypic similarity, we next considered the exon-intron structure of target genes. Specifically, we investigated phenotypes resulting from truncating mutations affecting the same exon in unrelated ASD probands; in this analysis, we took into account LGD mutations in the exon’s coding sequence as well as disruptions of the exon’s flanking canonical splice sites, since such splice site mutations should affect the same transcript isoforms (Supplementary Fig. 2). Interestingly, the analysis of 16 unrelated ASD probands (8 pairs with LGD mutations in the same exons) showed that they have strikingly more similar phenotypes (Fig. 1, red bars) compared to probands with LGD mutations in the same gene (Fig. 1, dark green bars); same exon FSIQ/NVIQ/VIQ average IQ difference 8.9, 8.3, 17.3 points, same gene average difference 28.3, 25.7, 34.9 points (Mann-Whitney U one-tail test *P* = 0.003, 0.005, 0.016). Because of well-known gender differences in autism susceptibility^11, 30, 31^, we also compared IQ differences between probands of the same gender harboring truncating mutations in the same exon (Fig. 1, orange bars) to IQ differences between probands of different genders; same gender FSIQ/NVIQ/VIQ average difference 5.4, 7.2, 12.2; different gender average difference 14.7, 10, 25.7 (MWU one-tail test *P* = 0.04, 0.29, 0.07). Thus, stratification by gender further decreases the phenotypic differences between probands with LGD mutations in the same exon. Notably, the patterns of phenotypic similarity between different probands only extended to mutations affecting the same exon. The average IQ differences between probands with LGD mutations in neighboring exons were not significantly different compared to mutations in non-neighboring exons (MWU one-tail test *P* = 0.6, 0.18, 0.8; Supplementary Fig. 3). The observed effects were also specific to LGD mutations; probands with either synonymous (*P* = 0.93, 0.97, 0.95; Supplementary Fig. 4) or missense (*P* = 0.8, 0.5, 0.8; Supplementary Fig. 5) mutations in the same exon were as phenotypically diverse as random pairs of ASD probands.

**Figure 1:**
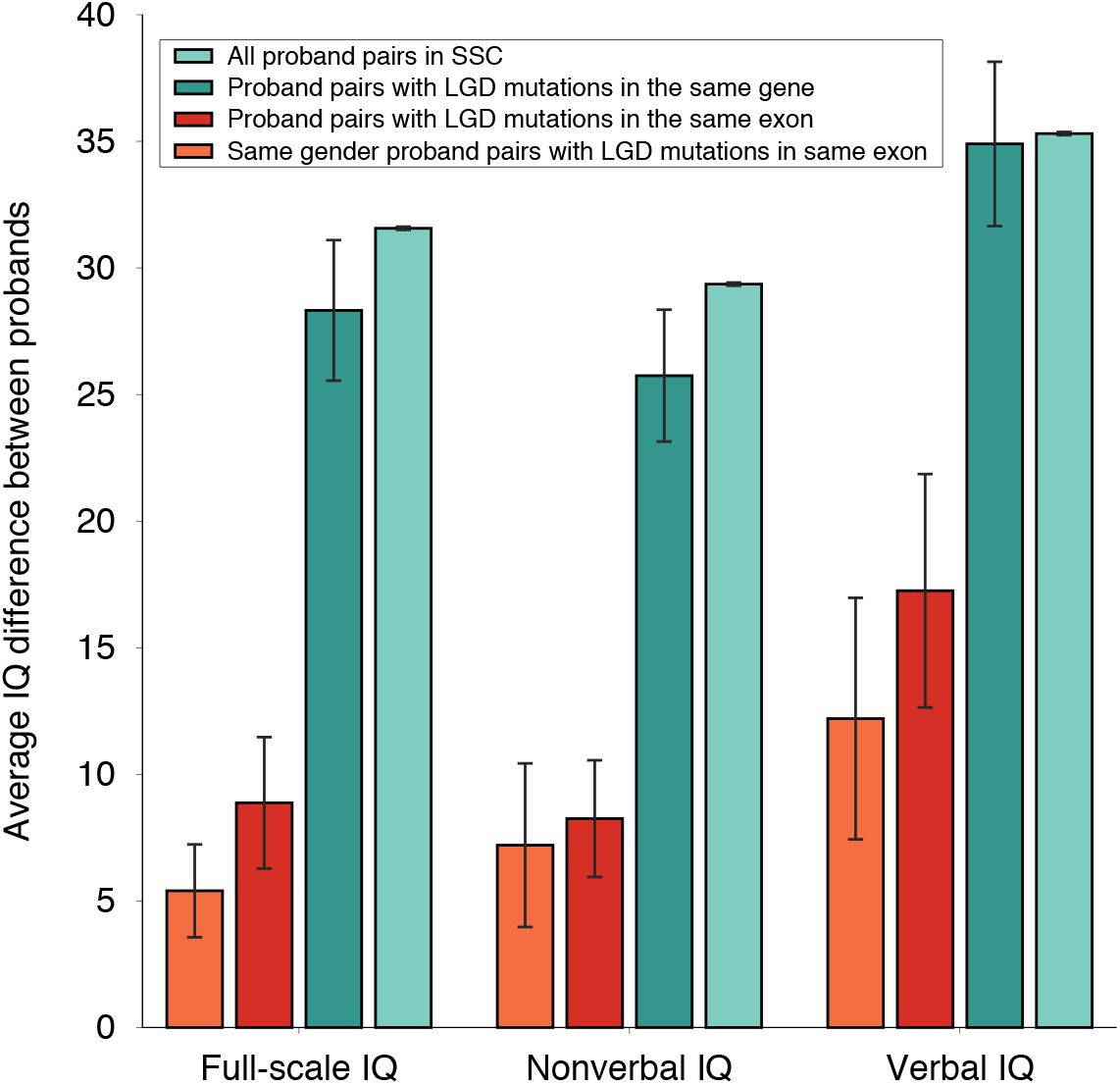
Average difference in IQs between SSC probands. From left to right, the sets of bars represent differences between full-scale, nonverbal, verbal IQs. Within each bar set, from right to left, the bars represent the average IQ difference between pairs of probands in the entire SSC cohort (light green), between probands with *de novo* LGD mutations in the same gene (dark green), between probands with *de novo* LGD mutations in the same exon (red), and between probands of the same gender and with *de novo* LGD mutations in the same exon (orange). Error bars represent the SEM.

We next explored the relationship between phenotypic similarity and the proximity of truncating mutations in the corresponding protein sequences. This analysis revealed that probands with LGD mutations in the same exon often had similar IQs, despite being affected by truncating mutations separated by scores to hundreds of amino acids in protein sequence (Fig. 2a; Supplementary Fig. 6). Furthermore, we found probands with LGD mutations in the same exon to be more phenotypically similar than probands with LGD mutations separated by comparable amino acid distances in the same protein sequence but not necessarily in the same exon (NVIQ distance-matched permutation test *P* = 0.002; Supplementary Fig. 7). We also investigated whether *de novo* mutations truncating a larger fraction of protein sequences resulted, on average, in more severe intellectual phenotypes. Surprisingly, this analysis showed no significant correlations between the fraction of truncated protein and the severity of intellectual phenotypes (Fig. 2b); NVIQ Pearson’s R = 0.05 (*P* = 0.35; Supplementary Fig. 8). We also did not find any significant biases in the distribution of truncating *de novo* mutations across protein sequences compared with the distribution of synonymous *de novo* mutations (Kolmogorov-Smirnov two-tail test *P* = 0.9; Supplementary Fig. 9). It is possible that the lack of the correlation between phenotypic impact and the fraction of truncated sequence is due to the averaging of various effects across different proteins. Therefore, for genes with recurrent mutations, we used a paired test to investigate whether truncating a larger fraction of the same protein sequence led to more severe phenotypes. This analysis also showed no substantial phenotypic difference due to LGD mutations truncating different fractions of the same protein (average NVIQ difference 0.24 points; Wilcoxon signed-ranked one-tail test *P* = 0.44). We also investigated, using the Pfam database^32^, whether mutations that truncate the same protein domain lead to more similar phenotypic differences. The results demonstrated that mutations in different exons, even when truncating the same protein domain, resulted, on average, in phenotypes as different as due to LGD mutations in the same protein (average NVIQ differences = 28.1; Supplementary Fig. 10).

**Figure 2:**
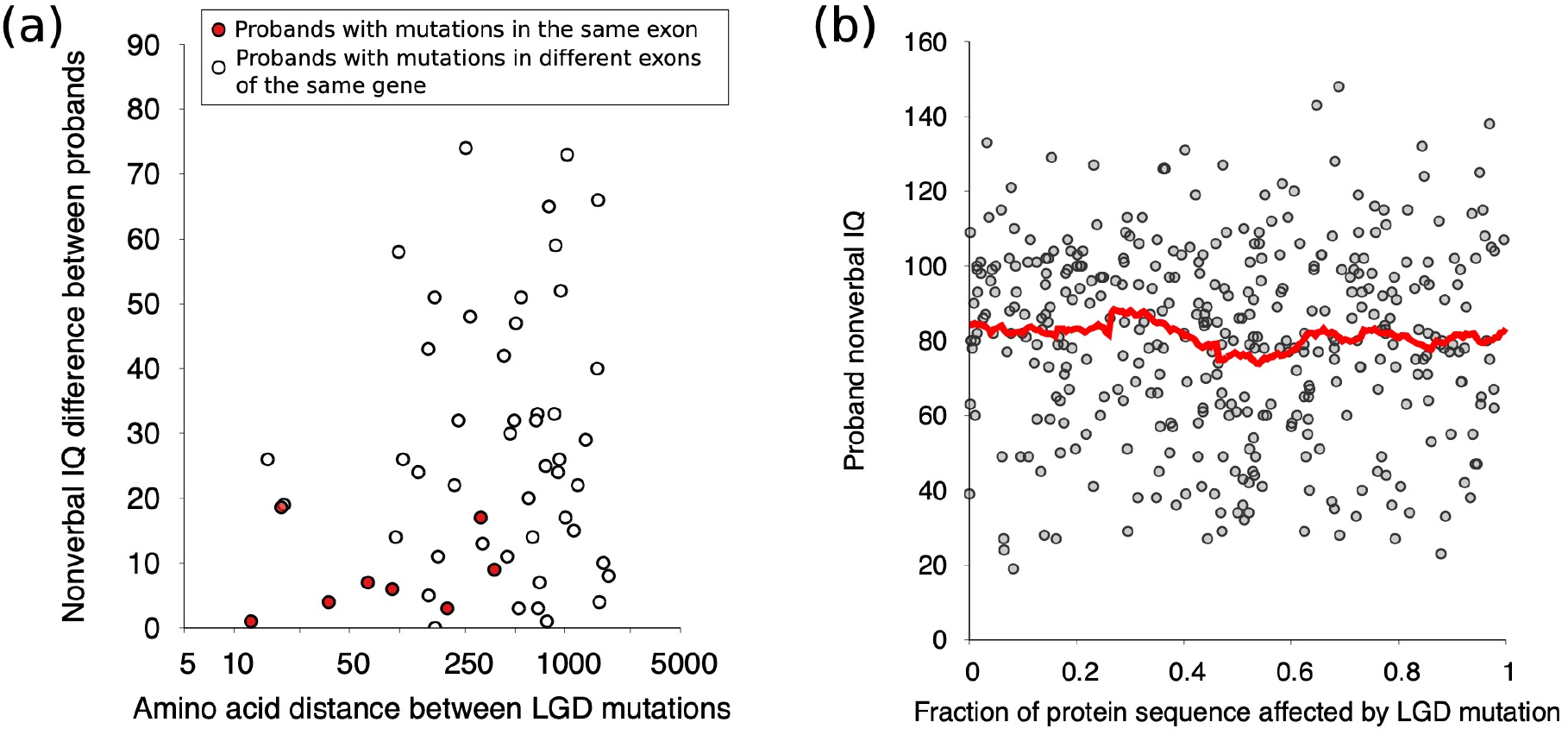
The relationship between the position of *de novo* LGD mutations in protein sequence and probands’ IQs. (**a**) Amino acid distance between LGD mutations in protein sequence versus differences in nonverbal IQ. Each point corresponds to a pair of probands with LGD mutations in the same gene. The x-axis represents the amino acid distance between LGD mutations, and the y-axis represents the difference between the corresponding probands’ nonverbal IQs (NVIQ). Red points represent pairs of probands with LGD mutations in the same exon, and white points represent pairs of probands with mutation in the same gene but different exons. (**b**) Relative fraction of protein sequence truncated by LGD mutations versus probands’ NVIQs. Each point corresponds to a single individual affected by an LGD mutation. The x-axis represents the fraction of protein sequence (i.e. fraction from the first amino acid) truncated by the mutation, and the y-axis represents the corresponding NVIQ. The red line represents a moving average of the data, calculated on an interval of width 0.05.

The results presented above suggest that the occurrence of *de novo* LGD mutations in the same exon, rather than simply the proximity of mutation sites in protein or nucleotide sequences, is primarily responsible for similar phenotypic consequences in unrelated probands. To explain this observation, we hypothesized that truncating mutations in the same exon usually affect, due to nonsense-mediated decay (NMD)^33^, the expression of exactly the same sets of splicing isoforms. Therefore, such mutations should lead to particularly similar phenotypes, both through similar decreases in overall gene dosage and similar perturbations to the mRNA expression of affected transcriptional isoforms. To evaluate this mechanistic model, we used data from the Genotype and Tissue Expression (GTEx) Consortium^34, 35^, which collected exome sequencing and corresponding human tissue-specific gene expression data from hundreds of individuals and across multiple tissues. Using ∼4,400 LGD variants in coding regions and corresponding RNA-seq data, we compared the expression changes resulting from LGD variants in the same and different exons of the same gene (Fig. 3a,b). Specifically, for each truncating variant, we analyzed allele-specific read counts^36^and then used an empirical Bayes approach to infer the effects of NMD on gene expression (see Methods). This analysis confirmed that the average gene dosage changes for individuals with LGD variants in the same exon were ∼7 times more similar compared to individuals with LGD variants in different exons of the same gene (Fig 3a); 2.2% versus 17.3% average difference in the decrease of overall gene dosage (Mann-Whitney U one-tail test *P* ˂ 2×10^−16^). Moreover, by analyzing GTEx data for each tissue separately, we found that across tissues LGD variants in the same exons lead to drastically more similar dosage changes of target genes (Fig. 3a).

**Figure 3:**
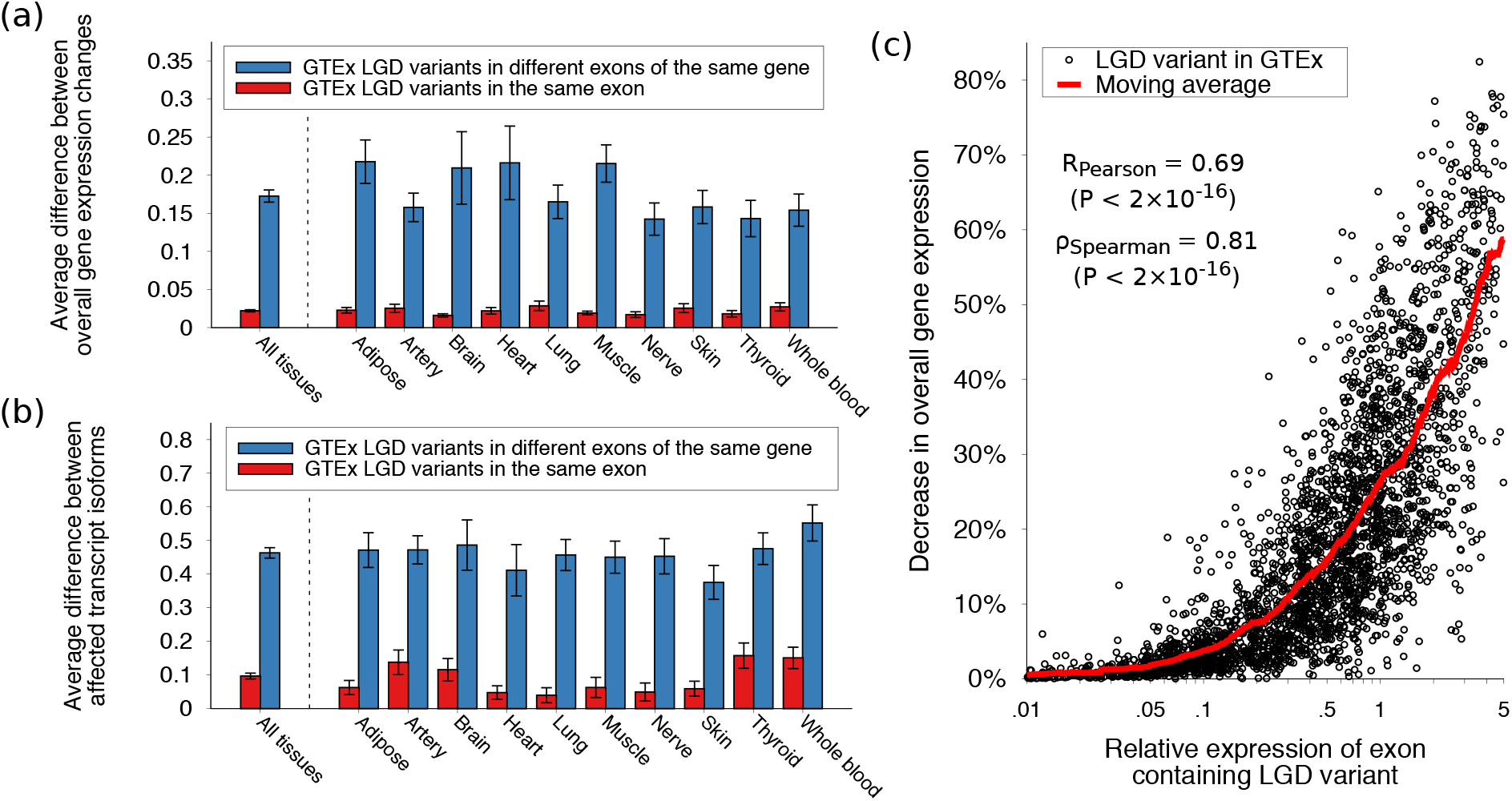
Gene expression changes across human tissues due to LGD variants in the same exon and in the same gene but different exons. Expression changes (decreases) due to LGD variants were calculated based on data from the Genotype and Tissue Expression (GTEx) Consortium^34^. (**a**) Bars represent the average difference across the GTEx cohort in overall gene expression changes induced by distinct LGD variants in the same exon (red) and in the same gene but different exons (blue). Error bars represent the SEM. (**b**) Bars represent the average difference across the GTEx cohort in isoform-specific expression changes induced by distinct LGD variants in the same exon (red) and in the same gene but different exons (blue). Differences in expression changes across transcriptional isoforms were quantified using the angular distance metric between vectors representing isoform-specific expression changes (see Methods). Error bars represent the SEM. (**c**) Relationship between the relative expression of exons containing LGD variants and the corresponding NMD-induced decreases in overall gene expression. Each point corresponds to an LGD variant in one of ten human tissues. The x-axis represents the relative expression of an exon harboring an LGD variant in a tissue; the relative expression of an exon was calculated as the ratio between the exon expression and total expression of the corresponding gene (see Methods). The y-axis represents the NMD-induced decrease in overall gene expression (see Methods). Red line represents a moving average of the data, calculated on an interval of width 0.1 (log-scaled).

Distinct splicing isoforms often have different functional properties^37, 38^. Consequently, LGD variants may affect phenotypes not only through NMD-induced changes in overall gene dosage, considered above, but also by altering the expression levels of different sets of splicing isoforms. To specifically analyze changes in the relative expression of distinct gene isoforms, we next used GTEx variants to quantify the effects of NMD on each isoform of a gene. To compare isoform-specific expression changes in the same gene, we calculated an angular distance metric between vectors representing dosage changes for each isoform (see Methods). This analysis demonstrated that changes in relative isoform expression are also significantly (∼5 fold) more similar for LGD variants in the same exon compared to variants in different exons of the same gene (Fig. 3b); 0.1 versus 0.46 for the average angular distance between isoform expression vectors (Mann-Whitney U one-tail test *P* < 2×10^−16^). These results were also consistent across tissues (Fig. 3b). Overall, the analyses of GTEx data demonstrate that both overall changes in gene dosage and changes in the relative expression levels of different isoforms are substantially more similar for truncating mutations in same exons.

Truncating variants in highly expressed exons should lead, on average, to relatively larger NMD-induced decreases in overall gene dosage. To confirm this hypothesis, we used RNA-seq data from GTEx. Specifically, for each exon harboring a truncating variant, we calculated its expression level relative to the expression values of the corresponding gene. We then explored the relationship between the relative expression of exons with the observed NMD-induced decreases in gene expression. The analysis indeed revealed a strong correlation between the relative expression levels of exons harboring LGD variants and the corresponding changes in overall gene dosage (Fig. 3c; Pearson’s R = 0.69, *P* < 2×10^−16^; Spearman’s ρ = 0.81, *P* < 2×10^−16^; see Methods). NMD-induced dosage changes may mediate the relationship between the relative expression levels of target exons and the corresponding phenotypic effects of truncating mutations. To investigate this relationship in detail, we used the BrainSpan dataset^39^, which contains exon-specific expression from human brain tissues. The BrainSpan data allowed us to estimate expression dosage changes resulting from LGD mutations in different exons of ASD-associated genes (see Methods).

Notably, it is likely that there is substantial variability across human genes in terms of the sensitivity of intellectual and other ASD phenotypes to gene dosage. Therefore, to quantify the sensitivity of IQ to changes in the expression of specific genes, we considered a simple linear dosage model. Specifically, we assumed for genes with recurrent truncating mutations in SSC that changes (decreases) in probands’ IQs are linearly proportional to the predicted relative decrease in overall gene dosage due to NMD. We further assumed that each human gene can be characterized by a parameter, which we call its phenotypic dosage sensitivity (PDS), characterizing the linear relationship between changes in gene dosage compared to wild type and the corresponding changes in a given human phenotype. Numerically, we defined IQ-associated PDS to be equal to the average change in IQ resulting from a 10% change in gene dosage. We restricted this analysis to LGD mutations predicted to cause NMD-induced expression changes, i.e. we excluded mutations within 50 bp of the last exon junction complex^40^, and also assumed the average neurotypical IQ (100) for wild type (intact) gene dosage. Using this linear model, for each gene with recurrent truncating ASD mutations, we used predicted changes in gene dosage to estimate gene-specific PDS parameters for intellectual phenotypes (Supplementary Fig. 11; see Methods). Notably, as we expected, PDS values varied substantially across 24 considered human genes (CV = SD/Mean = 0.57, for NVIQ).

We used the aforementioned linear model to explore the relationship between the relative expression values of exons (i.e. the ratio of exon expression to gene expression) harboring LGD mutations and the corresponding decreases in probands’ intellectual phenotypes. To account for differences in phenotypic sensitivity to dosage changes across genes, we normalized the observed changes in IQ by the estimated PDS values of affected genes. Normalized in this way, phenotypic effects represent changes in phenotype relative to the predicted effects for 10% decreases in dosage of affected genes. This analysis revealed that mutation-induced gene dosage changes are indeed strongly correlated with the normalized phenotypic effects; NVIQ Pearson’s R = 0.63, permutation test *P* = 0.02 (Fig. 4a; Supplementary Fig. 12). Very weak correlations were obtained for randomly permuted data, i.e. when truncating mutations were randomly re-assigned to different exons in the same gene (average NVIQ Pearson’s R = 0.18; see Methods). Since the heritability of intelligence is known to substantially increase with age^41^, we also investigated how the results depend on the age of probands. When we restricted our analysis to the older half of probands in SSC (i.e. older than the median age of 8.35 years), the strength of the correlations between the predicted dosage changes and normalized phenotypic effects increased further; NVIQ Pearson’s R = 0.75, average permuted R = 0.2, permutation test *P* = 0.019 (Fig. 4b; Supplementary Fig. 13). The strong correlations between target exon expression and intellectual ASD phenotypes suggest that, when gene-specific PDS values are taken into account, a significant fraction (30%-45%) of the relative phenotypic effects of *de novo* LGD mutations across genes can be explained by the resulting overall dosage changes of target genes.

**Figure 4:**
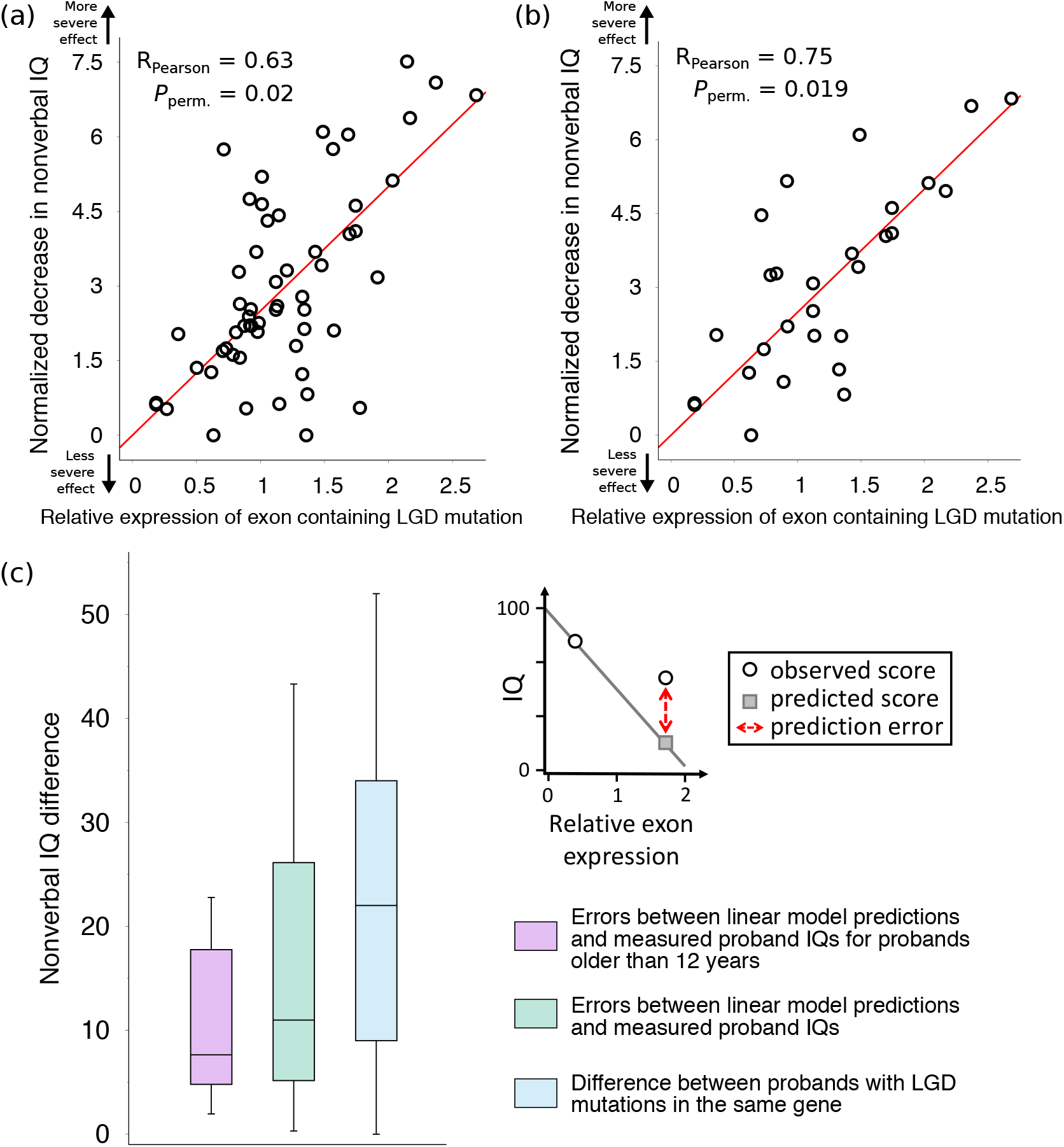
Relationship between the relative expression of exons harboring LGD mutations and the corresponding decrease in probands’ intellectual phenotypes. (**a**) Each point corresponds to a proband with an LGD mutation in a gene; only genes with multiple LGD mutations in the SSC cohort were considered. The x-axis represents the relative exon expression for exons harboring the LGD mutations. The y-axis represents the normalized decrease in the affected proband’s NVIQ, i.e. the NVIQ decrease divided by the NVIQ phenotypic dosage sensitivity (PDS) of that gene (see Methods). The regression line across all points is shown in red; *P*-values were calculated based on randomly shuffled data (see Methods). (**b**) Same as (a), but with the analysis restricted to the older half of SSC probands (i.e. older than the median age 8.35 years). (**c**) Boxplots represent the distribution of errors in predicting the effects of LGD mutations on NVIQ (see Methods); NVIQ prediction errors are shown for all probands (green), and for probands older than 12 years (purple). For comparison, the average differences in NVIQ scores between probands with LGD mutations in the same gene are also shown (blue). Only genes with multiple LGD mutations in SSC were considered. The ends of each solid box represent the upper and lower quartiles; the horizontal lines inside each box represent the medians; and the whiskers represent the 5^th^and 95^th^ percentiles. The inset panel illustrates the linear regression model used to perform leave-one-out predictions of probands’ NVIQs. Round open points represent observed phenotypic scores for probands with LGD mutations in the same gene, the grey square point represents the predicted phenotypic score, and the red dotted line represents the prediction error.

We next evaluated the ability of our linear dosage model, based on calculated PDS parameters, to explain the effects of LGD mutations on non-normalized IQs. To that end, for each gene with multiple truncating mutations in different probands, we used our linear regression model to perform leave-one-out predictions for IQ scores, i.e. we used PDS values calculated based on all but one probands with mutations in the gene to estimate IQ values for the left out proband (Fig. 4c, inset; see Methods). Despite the minimalism of our model and multiple simplifying assumptions, for LGD mutations that trigger NMD, the model median inference error for NVIQ was 11.1 points (Fig. 4c; Supplementary Fig. 14), which is significantly smaller than median NVIQ difference between probands with LGD mutations in the same gene, 22.0 points (MWU one-tail test *P* = 0.014). The NVIQ inferences based on probands of the same gender had significantly smaller errors compared to inferences based on probands of the opposite gender; same gender NVIQ median error 9.1 points, different gender median error 19.9 points (MWU one-tail test *P* = 0.018). Similar to normalized phenotypic effects (Fig. 4a,b), the inference errors decreased for older probands; for example, for probands older than 12 years, the median NVIQ inference error 7.6 points (Fig. 4c, Supplementary Fig. 14 and 15).

Given that relative exon usage varies across neural development^39, 42^, we investigated the relationship between developmental expression profiles of exons and ASD phenotypes. To that end, we sorted exons from genes harboring LGD mutations^14^into four groups (quartiles) based on their developmental expression bias, which was calculated as the fold-change between prenatal and postnatal exon expression levels (Fig. 5a). We then analyzed the enrichment of LGD mutations in each exon group (see Methods). Compared to exons with no substantial developmental bias, we found significant enrichment of LGD mutations not only in exons with a strong prenatal bias (binomial one-tail test *P* = 8×10^3^, Relative Rate = 1.33), but also in exons with postnatal biases (*P* = 0.018, RR = 1.31) (Fig. 5b). To understand the origin of the observed exon biases, we stratified probands into lower (≤ 70) and higher IQ (> 70) cohorts (Fig. 5c). Interestingly, while LGD mutations associated with lower IQs were strongly enriched only in prenatally biased exons (binomial one-tail test *P* = 6×10^−3^, RR = 1.62), mutations associated with higher IQs were exclusively enriched in postnatally biased exons (*P* = 0.05, RR = 1.27). These results demonstrate that mutations in exons with biases towards prenatal and postnatal expression preferentially contribute to ASD cases with lower and higher IQ phenotypes, respectively. We note that the observed exon developmental biases for LGD mutations are not simply driven by biases at the gene level, as mutations associated with both higher and lower IQ phenotypes showed enrichment exclusively towards genes with prenatally biased expression (Supplementary Fig. 16).

**Figure 5:**
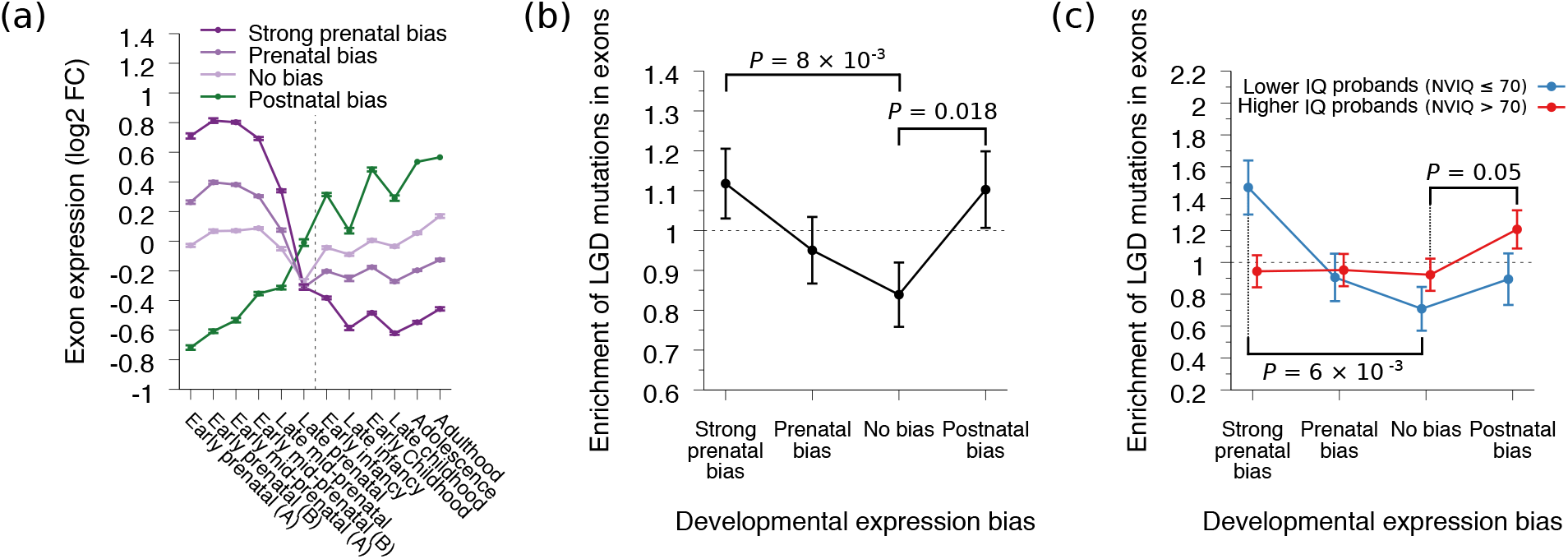
Relationship between the developmental expression of exons and intellectual ASD phenotypes. (**a**) Exon developmental expression profiles for genes with *de novo* LGD mutations in SSC. Exons from all genes harboring LGD mutations were sorted into four groups (“strong prenatal bias”, “prenatal bias”, “no bias”, and “postnatal bias”) based on their overall developmental expression bias; the developmental bias was calculated as the log2 fold change between the average prenatal and postnatal exon expression levels. Lines represent the average expression profiles for exons in each group, and the x-axis represents 12 periods of human brain development, based on data from the Allen Institute’s BrainSpan atlas ^39^. The vertical dotted line delineates prenatal and postnatal developmental periods. Error bars represent the SEM. (**b**,**c**) Enrichment of LGD mutations across the four exon groups with different developmental biases. The y-axes represent the enrichment (relative rate) of mutations in each exon group; the enrichment was calculated by randomizing LGD mutations across exons proportionally to the exons’ coding sequence lengths (see Methods). Error bars represent the SEM. (**b**) The overall enrichment of LGD mutations across the four exon groups of exons with different developmental expression biases. (**c**) The enrichment of LGD mutations across the four exon groups calculated separately for ASD probands with higher (>70, red) and lower (≤70, blue) nonverbal IQs.

Although we primarily analyzed the impact of autism mutations on intellectual phenotypes, similar dosage and isoform expression changes of affected genes may also lead to analogous results for other quantitative ASD phenotypes^24, 43^. Indeed, for LGD mutations predicted to lead to NMD, we observed similar patterns for several other key autism phenotypes. Specifically, SSC probands with truncating mutations in the same exon exhibited more similar adaptive behavior abilities compared to probands with mutations in the same gene (Fig. 6a, left set of bars, Supplementary Fig. 17); Vineland Adaptive Behavior Scales (VABS)^44^composite standard score difference of 4.7 versus 12.1 points (Mann-Whitney U one-tail test *P* = 0.017). In contrast, VABS differences between probands with truncating mutations in the same gene were not significantly different than for randomly paired probands (Fig. 6a, Supplementary Fig. 17); 12.1 versus 13.7 points (MWU one-tail test *P* = 0.23). Furthermore, probands with truncating mutations in the same exon also displayed more similar fine motor skills; in the Purdue Pegboard Test, 1.2 versus 3.0 for the average difference in normalized tasks completed with both hands (MWU one-tail test *P* = 0.02; Supplementary Fig. 18; see Methods). Coordination scores in the Social Responsiveness Scale questionnaire were also more similar in probands with LGD mutations in the same exon compared to probands with mutations in the same gene; 0.6 versus 1.1 for the average difference in normalized response (MWU one-tail test *P* = 0.05; Supplementary Fig. 19).

**Figure 6:**
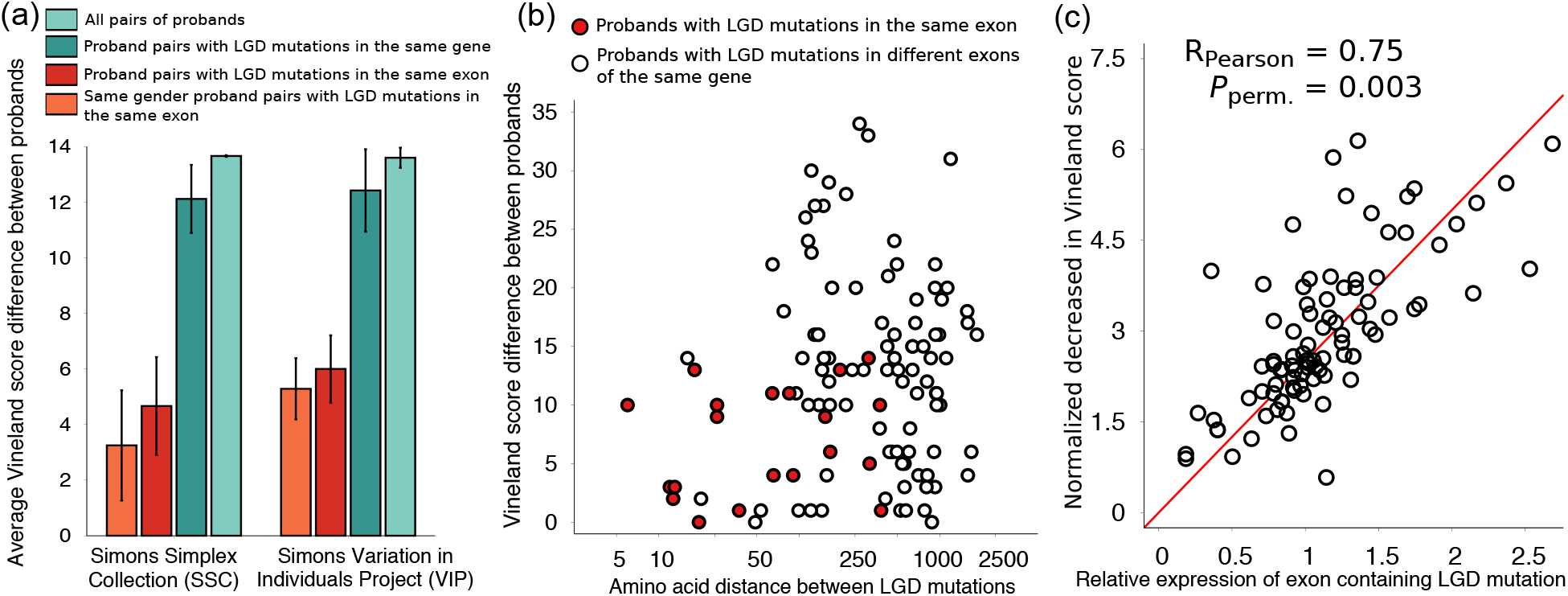
Validation of the observed IQ patterns using Vineland Adaptive Behavior Scales (VABS) scores. (**a**) Average VABS score differences between probands using data from SSC (left set of bars) and VIP (right set of bars). Each bar shows the average difference in VABS scores between pairs of probands in different groups. In the SSC and VIP bar sets, from right to left, bars represent differences between all pairs of probands in each cohort (light green), between probands with LGD mutations in the same gene (dark green), between probands with LGD mutations in the same exon (red), and between probands of the same gender with LGD mutations in the same exon (orange). Error bars represent the SEM. (**b**) Amino acid distance between LGD mutations in the same protein versus differences in VABS score. Each point corresponds to a pair of probands, with individual from either SSC or VIP, with LGD mutations in the same gene. The x-axis represents the amino acid distance between LGD mutations, and the y-axis represents the difference between the affected probands’ VABS scores. Red points represent proband pairs with LGD mutations in the same exon, and white points represent proband pairs with LGD mutations in different exons of the same gene. (**c**) Relationship between the relative expression of exons harboring LGD mutations and the corresponding decrease in probands’ normalized VABS scores. Each point corresponds to a proband with an LGD mutation in a gene; only genes with multiple LGD mutations were considered. The x-axis represents the relative expression (exon expression divided by total gene expression) of exons harboring LGD mutations. The y-axis represents the affected probands’ normalized decrease in VABS scores, i.e. the VABS decrease divided by the VABS phenotypic dosage sensitivity (PDS) of that gene. The regression line across all points is shown in red; *P*-values were calculated based on randomly shuffled data (see Methods). The analysis was restricted to *de novo* LGD mutations predicted to trigger NMD, i.e. more than 50 bp upstream from the last exon junction.

Finally, we sought to validate the observed phenotypic patterns using an independent cohort of ASD probands. To that end, we analyzed an independently collected dataset from the ongoing Simons Variation in Individuals Project (VIP)^29^. The analyzed VIP dataset contained genetic information and VABS phenotypic scores for 41 individuals with *de novo* LGD mutations in 12 genes. Reassuringly, and consistent with our findings in SSC, probands from the VIP cohort with truncating *de novo* mutations in the same exon also exhibited strikingly more similar VABS phenotypic scores compared to probands with mutations in the same gene (Fig. 6a, right set of bars, Supplementary Fig. 20); VABS composite standard score difference 6.0 versus 12.4 (Mann-Whitney U one-tail test *P* = 0.014). Similar to the SSC cohort, LGD mutations in neighboring exons did not result in more similar behavior phenotypes; VABS composite standard score average difference 13.6 points (MWU one-tail test *P* = 0.6). The fraction of truncated proteins also did not show significant correlation with the VABS scores of affected probands (Pearson’s R = −0.08, *P* = 0.7). Using VABS scores from both SSC and VIP, we next investigated whether, analogous to the IQ phenotypes (Fig. 3a), the similarity of VABS scores is primarily due to the presence of mutations in the same exon, rather than proximity of truncating mutations within the corresponding protein sequence. Indeed, LGD mutations in the same exon often resulted in similar adaptive behavior abilities even when the corresponding mutations were separated by hundreds of amino acids (Fig. 6b, red points, Supplementary Fig. 21). By comparing mutations in the same exon to mutations separated by similar amino acid distances in the same protein but not necessarily the same exon, we confirmed that probands with mutations in the same exon were significantly more phenotypically similar (permutation test *P* = 3×10^−4^; Supplementary Fig. 22; see Methods). When we applied the linear dosage model while accounting for the VABS sensitivity to changes in the dosage of different genes (i.e. gene-specific PDS values), we found substantial correlations between the relative expression of exons harboring LGD mutations and the normalized VABS phenotypes of the affected probands (Pearson R = 0.75, permutation test *P* = 0.003; Fig. 6c; Supplementary Fig. 23). Overall, these results confirm the generality of the phenotypic patterns observed in the SSC cohort.

## Discussion

Previous studies explored phenotypic similarity in syndromic forms of ASD due to mutations in specific genes^(45–49^. Nevertheless, across a large collection of contributing genes, the nature of the substantial phenotypic heterogeneity in ASD is not well understood. Interestingly, the diversity of intellectual and other important ASD phenotypes resulting from *de novo* LGD mutations in the same genes is usually only slightly (∼10%) smaller than the phenotypic diversity across the entire ASD cohort (Fig. 1, Fig. 6a). The presented results suggest that truncating mutations usually result in a range of relatively mild NMD-induced gene dosage changes, on average decreasing gene expression by ∼15-30% (Supplementary Fig. 24; see Methods). Our study further suggests a hierarchy of biological mechanisms contributing to phenotypic heterogeneity in simplex ASD cases triggered by LGD mutations in different genes and within the same gene.

Across LGD mutations, there is a significant but small correlation between a target gene’s brain expression level and the resulting intellectual phenotype (R^2^= 0.02, *P* = 0.03). This correlation is small, at least in part, due to the significant variability of expression levels across different exons in a gene. Indeed, intellectual phenotypes correlate significantly better with the relative expression level of exons harboring LGD mutations (R^2^= 0.10, *P* = 0.011). In addition to effects associated with different expression levels of exons, there is also substantial variability in the sensitivity of each specific phenotype to dosage changes of different genes. When we account for varying dosage sensitivities using gene-specific PDS values, the correlation between predicted dosage changes and normalized phenotypic effects becomes substantial (R^2^=0.4, *P* = 0.02, Fig. 4a; Fig. 6c). As the heritability of IQ phenotypes usually increases with age, we observe even stronger dosage-phenotype correlations for older probands (R^2^= 0.56, *P* = 0.019, Fig. 4b). Furthermore, even perturbations leading to similar dosage changes in the same gene may result in diverse phenotypes in cases where different, functionally distinct splicing isoforms are truncated. However, when exactly the same sets of isoforms are perturbed, as for LGD mutations in the same exon, the resulting phenotypes, even in unrelated ASD probands, become especially similar (Fig. 1, Fig. 6a). For LGD mutations affecting intellectual phenotypes, we found that same exon membership accounts for a larger fraction of phenotypic variance than multiple other genomic features, including expression, evolutionary conservation, pathway membership, and domain truncation (see Methods). There are likely deviations from the aforementioned patterns for specific genes and specific truncating mutations. For example, truncated proteins that escape NMD may lead to partial buffering, due to remaining activity, or to further damaging effects, due to dominant negative interactions. Nevertheless, our results demonstrate that for *de novo* LGD mutations in ASD, exons, rather than genes, usually represent a unit of effective phenotypic impact.

Our results also suggest that changes in ASD phenotypes induced by LGD mutations may be characterized by a simple linear model quantifying the sensitivity of a phenotype to changes in gene dosage. We observe that PDS values for the same phenotype vary substantially across genes (Supplementary Fig. 11), and that PDS differences are a major source of phenotypic variability. Moreover, PDS values for the same gene vary across phenotypes (for example, correlation between PDS values for IQ and VABS across 24 genes, R^2^= 0.37, P-values=0.001), which suggests that PDS values are specific to phenotype-gene pairs. Although we evaluated PDS parameters using predicted NMD-induced dosage changes, it may be possible to infer these parameters using other mechanisms of dosage change, such as regulatory mutations. As genetic and phenotypic data accumulate, it will be interesting to estimate gene-specific PDS values for multiple phenotypes and for a substantial number of ASD risk genes. Furthermore, due to consistent patterns of gene expression changes across tissues (Fig. 3), it may be possible to estimate PDS parameters for other genetic disorders and phenotypes. We note in this respect that quantitative gene-dosage relationships have been recently characterized for yeast fitness values in different environmental conditions^50^.

In the present study, we focused specifically on simplex cases of ASD, in which *de novo* LGD mutations are highly penetrant and where the contribution of genetic background is minimized. It is likely that differences in genetic background and environment represent other important sources of phenotypic variability^22, 51, 52^. Therefore, in more diverse cohorts, individuals with LGD mutations in the same exon will likely display greater phenotypic heterogeneity. For example, the Simons Variation in Individuals Project identified broad spectra of phenotypes associated with specific variants in the general population^29, 53–55^. We also observed significantly larger phenotypic variability for probands from sequenced family trios, i.e. families without unaffected siblings (Supplementary Fig. 25). For these probands, the enrichment of *de novo* LGD mutations is substantially lower and the contribution from genetic background is likely to be larger^56^, resulting in more pronounced phenotypic variability.

Our study may have important implications for precision medicine^51, 57, 58^. The presented results indicate that relatively mild decreases in affected gene dosage may account for a substantial fraction of adverse phenotypic consequences. Thus, from a therapeutic perspective, compensatory expression of intact alleles, as was recently demonstrated in mouse models of ASD^(59–61^and other diseases^62^, may provide an approach for alleviating phenotypic effects for at least a fraction of ASD cases. From a prognostic perspective, our results suggest that by sequencing and phenotyping sufficiently large patient cohorts with truncating mutations in different exons, it may be possible to understand likely phenotypic consequences originating from LGD mutations in specific exons. Furthermore, because we observe consistent patterns of expression changes across multiple human tissues, similar analyses may be also extended to other disorders affected by highly penetrant truncating mutations.

## Acknowledgements

We thank Drs. W.K. Chung, I. Pe’er, A. Packer, and all members of the Vitkup lab for helpful scientific discussions. D.V. acknowledges funding from the Simons Foundation (SFARI #308962). This work was supported in part by NIH grant no. T15LM007079 (A.H.C., J.C., J.W.) and Ruth L. Kirschstein National Research Service Award Institutional Research Training grant no. T32GM082797 (A.H.C.).

## Conflict of Interest statement

The authors declare that there is no conflict of interest.

## Notes

### Competing Interest Statement

The authors have declared no competing interest.

### Summary of Updates

Manuscript revised with additional background/introductory text. Abstract revised. Figures reformatted for clarity.

