## Supplementary material for "Exons as units of phenotypic impact for truncating mutations in autism": Supllementary Figures

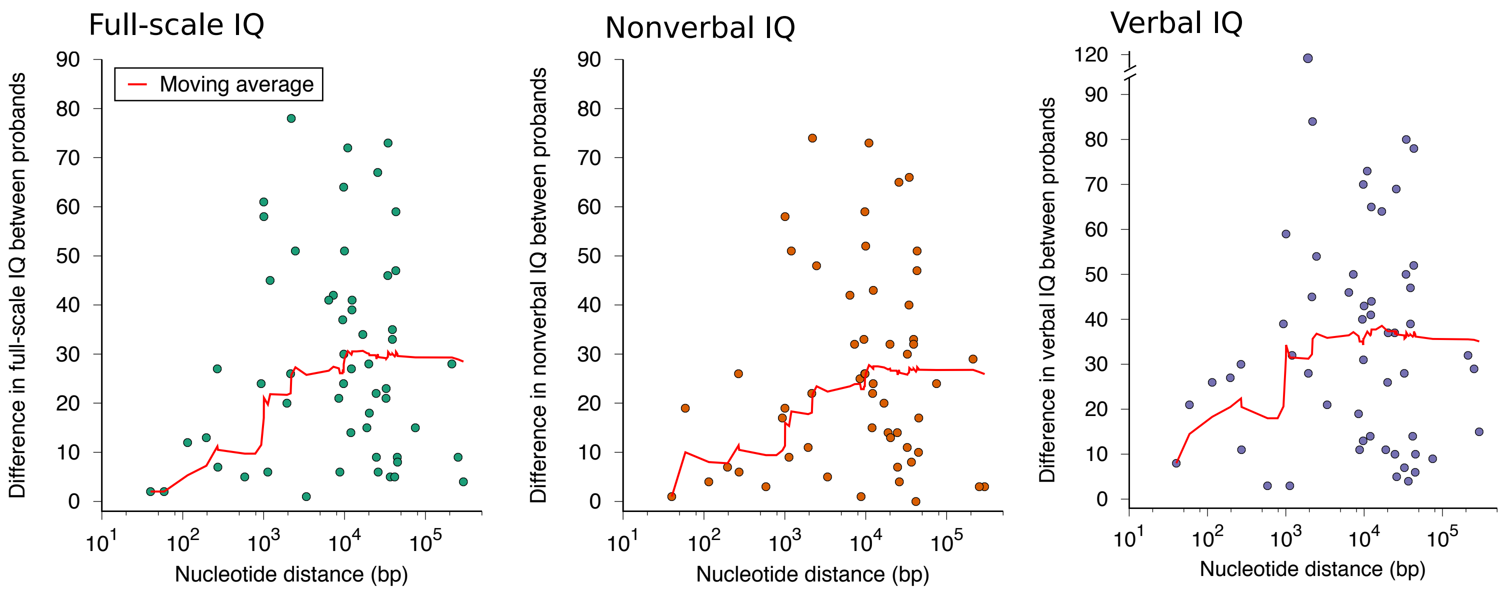


**Supplementary Figure 1**: IQ differences between pairs of probands with *de novo* LGD mutations in the same gene. Each point in the figures corresponds to a pair of probands from the SSC cohort with *de novo* LGD mutations in the same gene. The x-axis represents the nucleotide distance between LGD mutations. The y-axis represents the absolute difference in IQs (full-scale, nonverbal, or verbal IQ) between affected probands. Moving averages are shown in red.

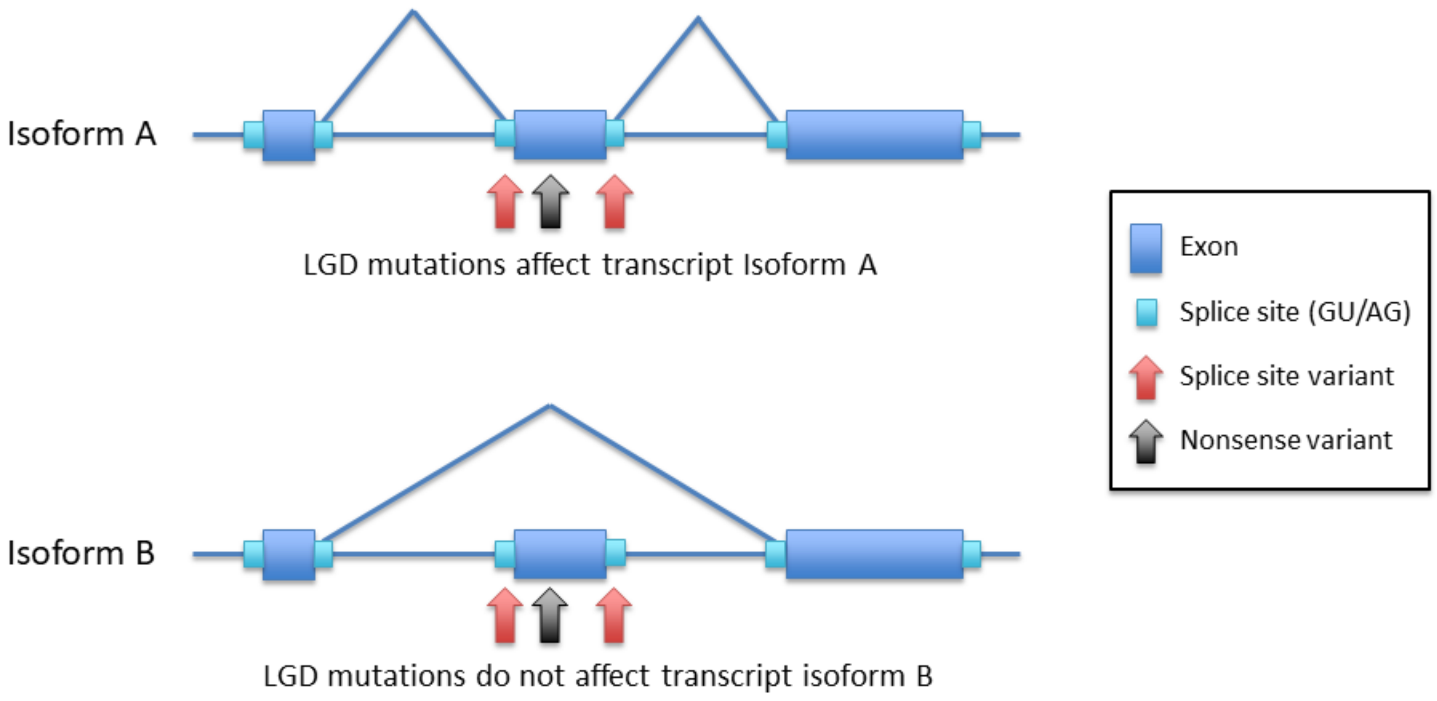


**Supplementary Figure 2:** Illustration showing an example of LGD mutations affecting either of two transcriptional isoforms of a gene. Exons are represented by blue rectangles, their flanking canonical splice sites by light blue boxes, and splicing patterns by diagonal lines joining the corresponding splice sites. Loss-of-function mutations in the exon's coding sequence (black arrows) and mutations disrupting the exon's flanking canonical splice sites (red arrows) usually affect the same transcriptional isoforms.

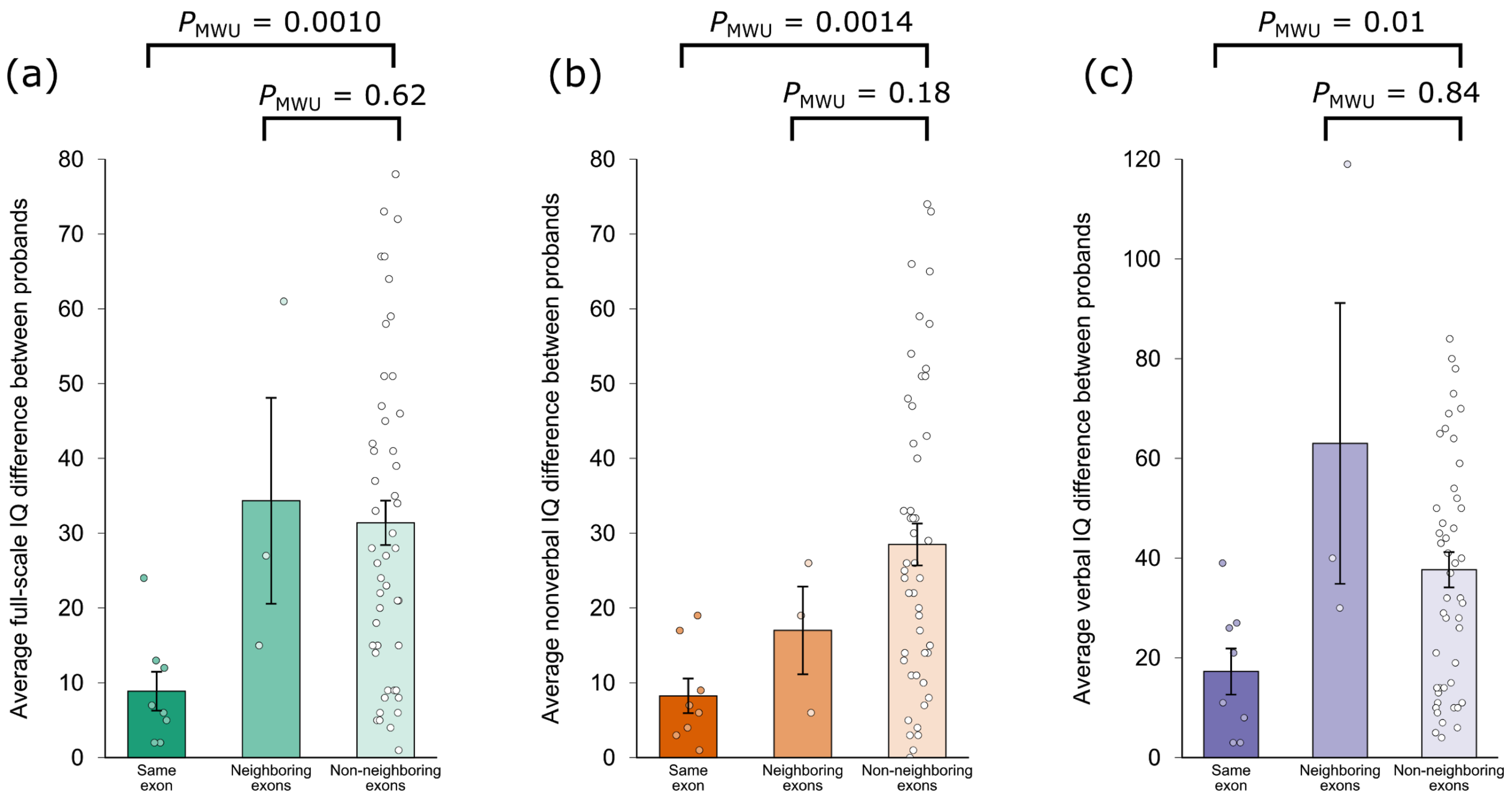


**Supplementary Figure 3:** The average IQ differences between probands with LGD mutations in the same exon, in neighboring exons of the same gene, and in non-neighboring exons of the same gene. Plots represent **(a)** full-scale IQ (FSIQ), **(b)** nonverbal IQ (NVIQ), and **(c)** verbal IQ (VIQ) scores. Each overlaid point represents a pair of probands with LGD mutations in the same exon, in neighboring exons, or in non-neighboring exons. The y-axis represents the IQ difference between affected probands. The statistical significance was determined using Mann-Whitney U tests ($P_{MWU}$). Error bars represent the SEM.

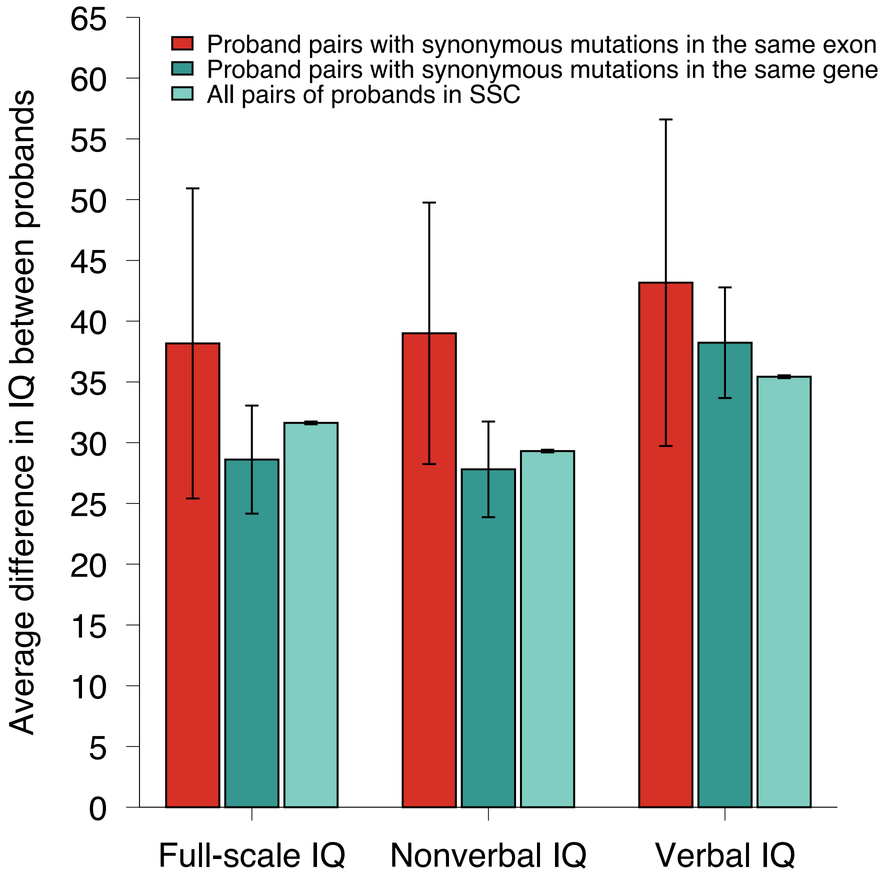


**Supplementary Figure 4**: The average differences in IQ between probands with synonymous mutations in the same gene or the same exon. Each bar represents the average IQ difference between all pairs of probands in the SSC cohort (light green), between pairs of probands with synonymous mutations in the same gene (dark green), and between pairs of probands with synonymous mutations in the same exon (red). From left to right, sets of bars represent differences for full-scale, nonverbal, and verbal IQs. Error bars represent the SEM.

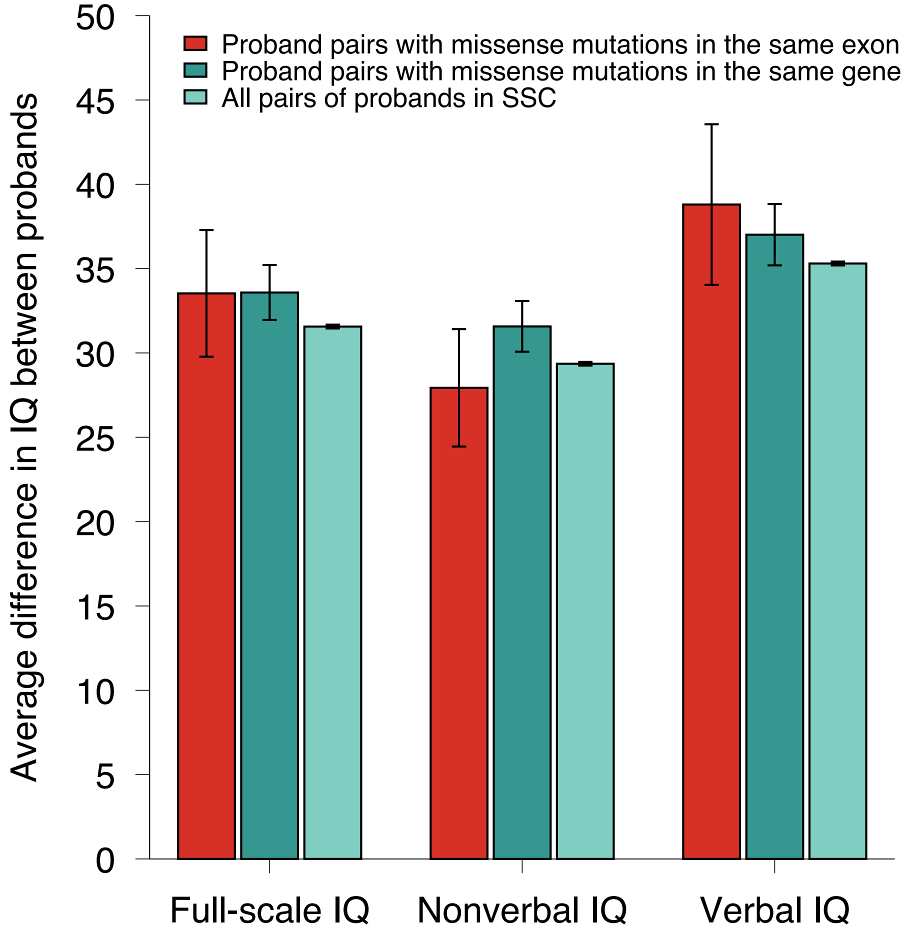


**Supplementary Figure 5:** Average difference in IQ between probands with missense mutations in the same gene or the same exon. Each bar represents the average IQ difference between all pairs of probands in the SSC cohort (light green), between pairs of probands with missense mutations in the same gene (dark green), and between missense mutations in the same exon (red). Error bars represent the SEM.

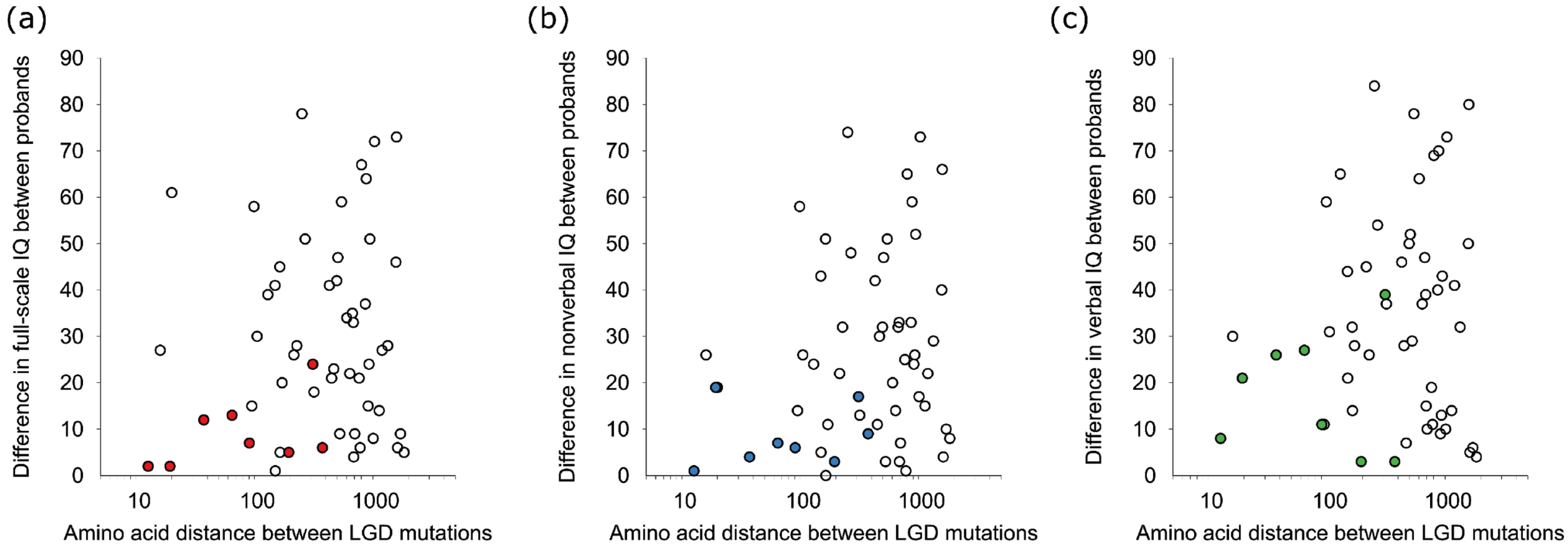


**Supplementary Figure 6:** Amino acid distance between LGD mutations in protein sequence versus the IQ differences between corresponding probands. From left to right, plots show differences in **(a)** full-scale IQ (FSIQ), **(b)** nonverbal IQ (NVIQ), and **(c)** verbal IQ (VIQ) scores. Each point in the figures corresponds to a pair of probands affected by *de novo* LGD mutations in the same gene. The x-axis represents the amino acid distance between LGD mutations, and the y-axis represents the absolute difference between the probands' IQs. Open (white) points correspond to pairs of probands with LGD mutations in different exons of the same genes; filled (colored) points correspond to pairs of probands with mutations in the same exon of the same genes.

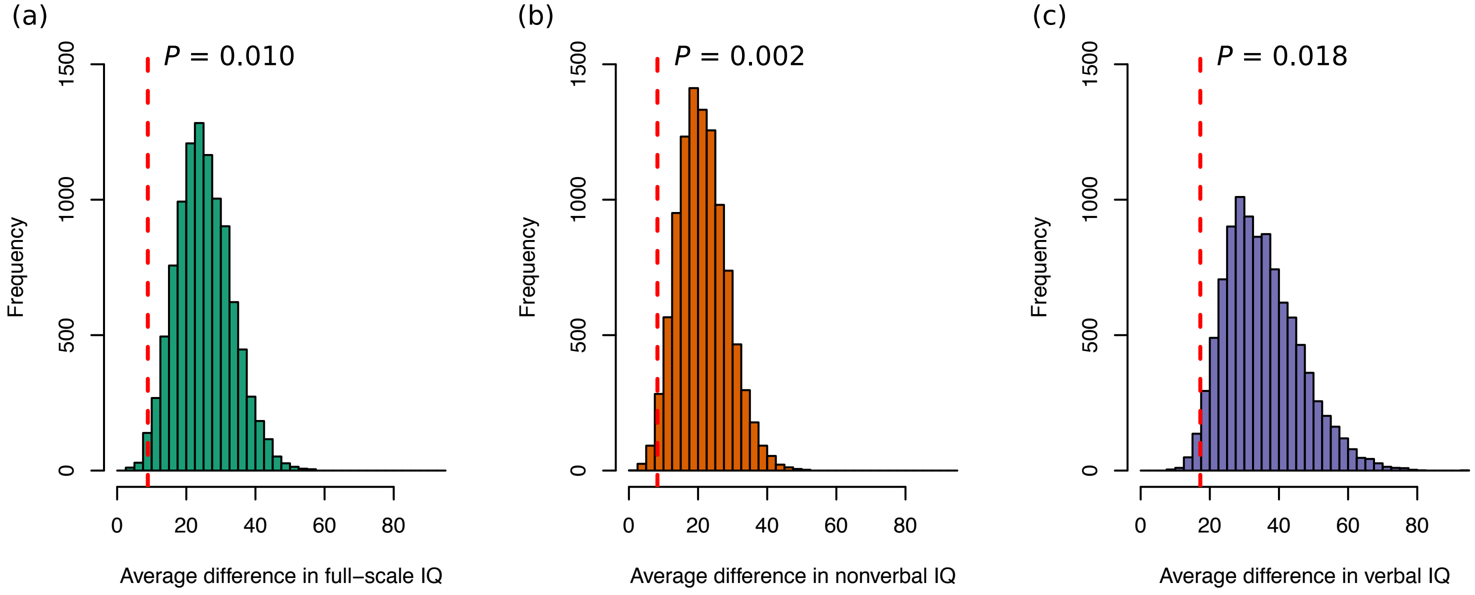


**Supplementary Figure 7:** Simulated null distributions of the average IQ difference between probands with *de novo* LGD mutations in the same gene and with similar distances in the corresponding protein sequence. From left to right, histograms show the distribution of average differences for **(a)** full-scale IQ (FSIQ), **(b)** nonverbal IQ (NVIQ), and **(c)** verbal IQ (VIQ) scores across the sampled null distribution trials (N=10,000). The histograms show the null trial frequencies of the average IQ differences (x-axes) between pairs of probands with mutations in the same gene, where mutations were separated by amino acid distances similar to the ones empirically observed between LGD mutations in the same exon (see Methods). The red dashed lines represent the observed average IQ differences between probands with LGD mutations in the same exon.

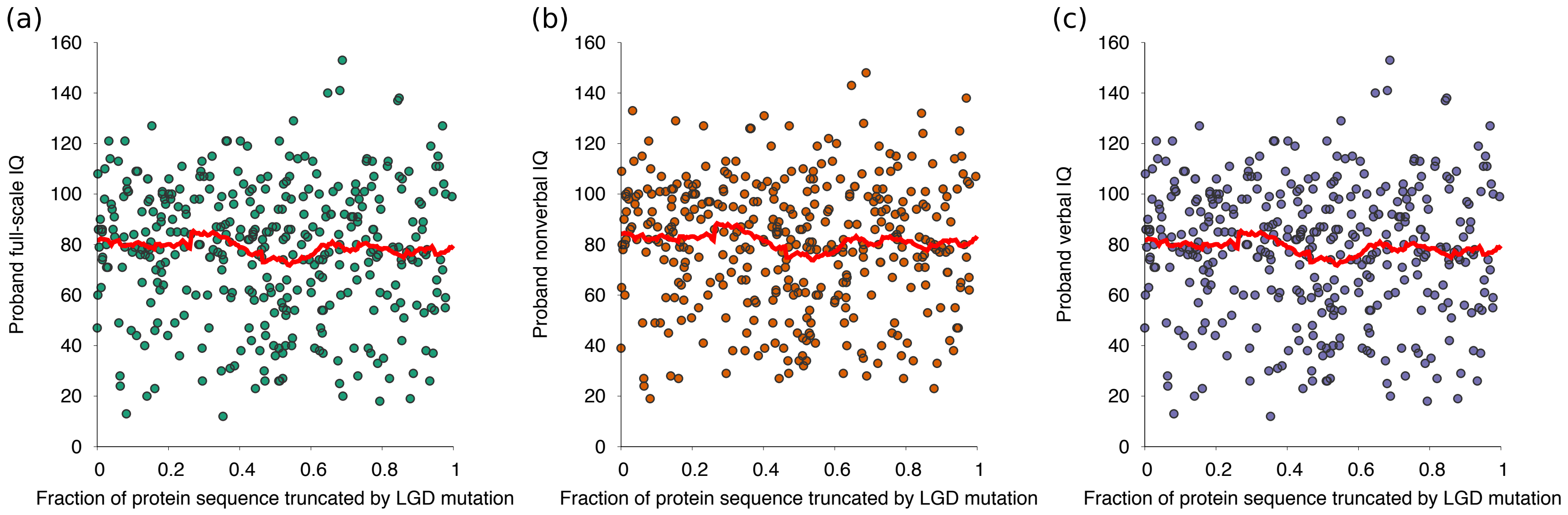


**Supplementary Figure 8:** Relative fraction of protein sequence truncated by LGD mutations versus proband IQs. Each point corresponds to a single proband in SSC affected by an LGD mutation. From left to right, the plots show **(a)** full-scale IQ (FSIQ), **(b)** nonverbal IQ (NVIQ), and **(c)** verbal IQ (VIQ) scores. The x-axis represents the fraction of protein amino acid sequence (i.e. fraction from the first amino acid) truncated by the LGD mutation. The y-axis represents the corresponding proband's IQ score. Red lines represent moving averages of the data.

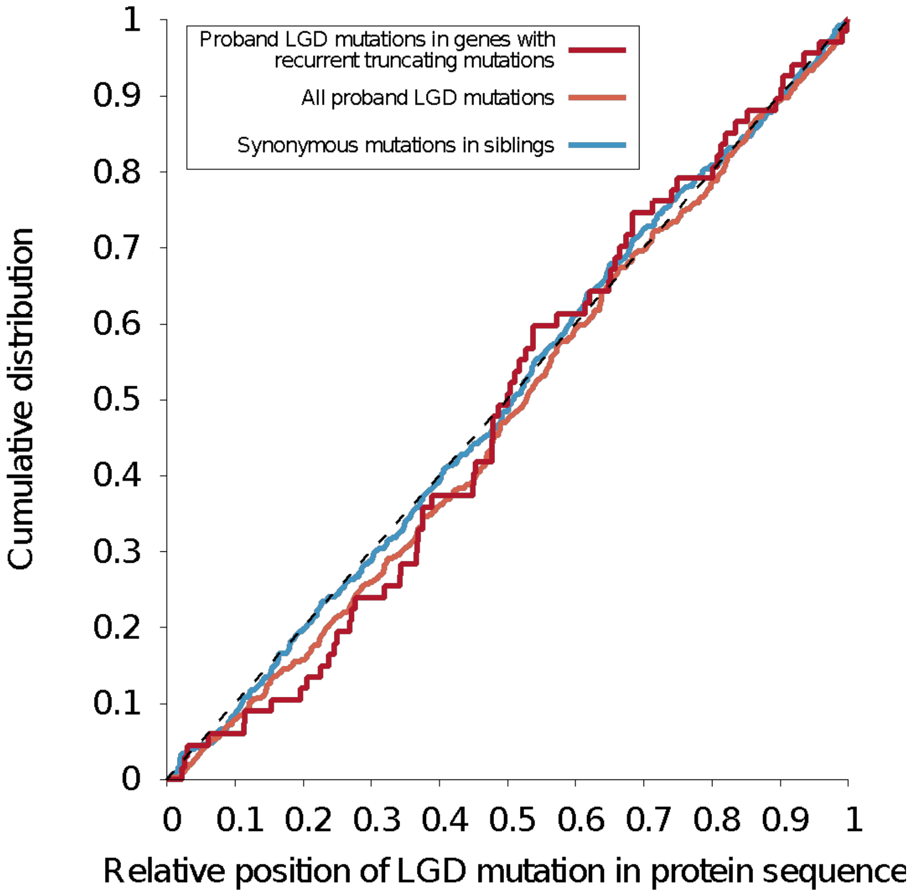


**Supplementary Figure 9:** Cumulative distributions of the relative amino acid positions of *de novo* LGD mutations. Each line represents the cumulative distribution of the relative protein sequence positions (i.e. fraction of the total sequence length from the first amino acid) for LGD mutations in genes with multiple truncating mutations in SSC (red), for all LGD mutations in SSC (orange), and for synonymous mutations in unaffected siblings in SSC (blue).

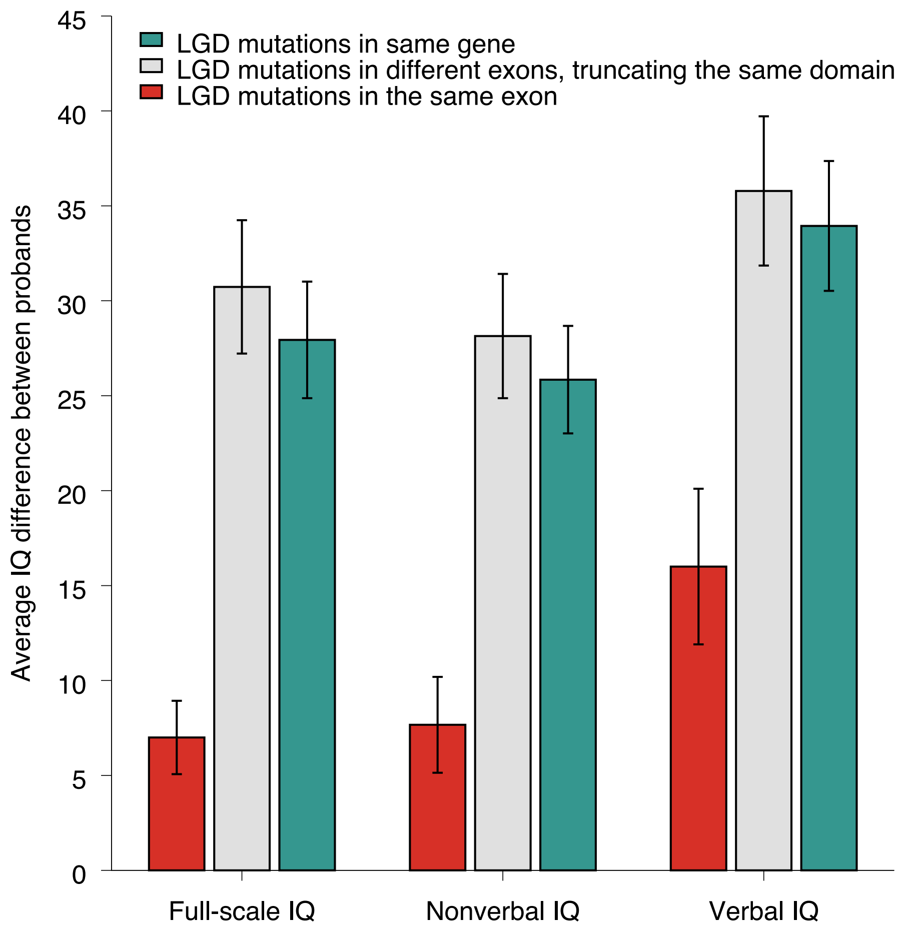


**Supplementary Figure 10:** Average phenotype difference between probands with LGD mutations truncating the same protein domain for full-scale, nonverbal, and verbal IQs. Bars represent the average difference in phenotypic scores between pairs of probands with LGD mutations in the same gene (dark green), in different exons that truncate the same protein domain (grey), and in the same exon (red). Error bars represent the SEM.

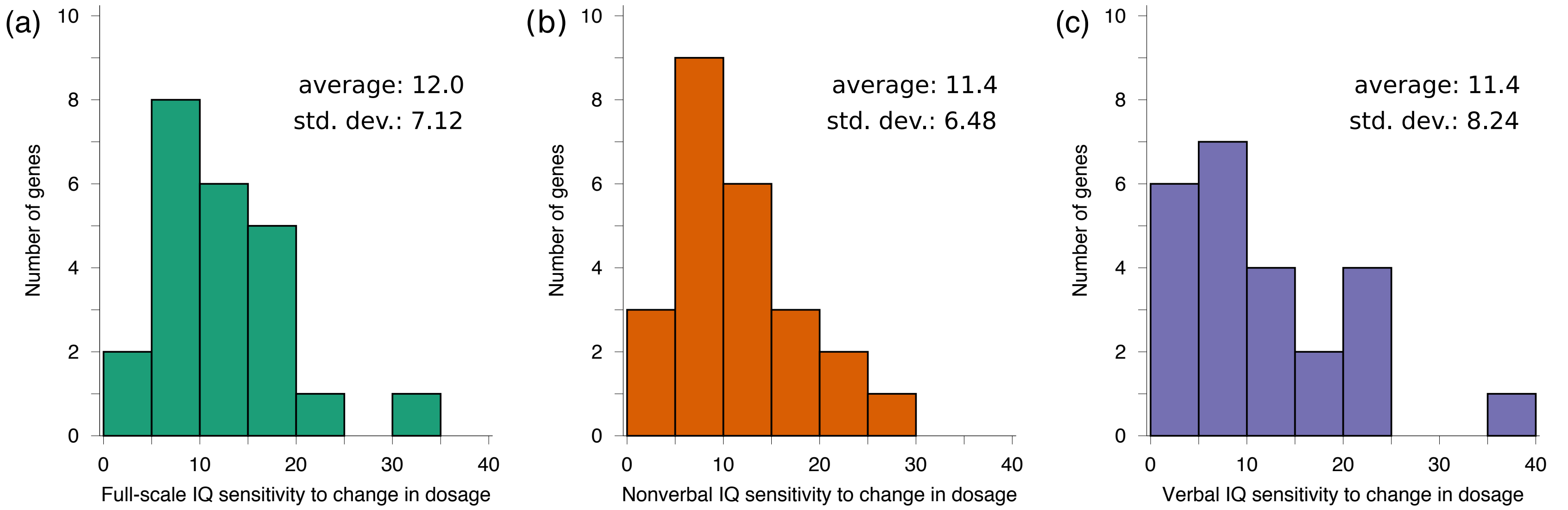


**Supplementary Figure 11:** Distribution sensitivity of the IQ phenotype to changes in gene dosage, i.e. the Phenotype Dosage Sensitivity (PDS), across different ASD genes with recurrent truncating mutations in SSC. From left to right, the histograms show the distribution of PDS sensitivity parameters across genes for **(a)** full-scale IQ (FSIQ), **(b)** nonverbal IQ (NVIQ), and **(c)** verbal IQ (VIQ). To calculate PDS parameters, we used linear regression, separately for each gene, to fit the relationship between the observed IQ decrease, compared to the neurotypical average value (100), and the estimated change in gene dosage due to LGD mutations (see Methods). We then calculated PDS as the predicted decrease in IQ due to 10% decrease in gene dosage.

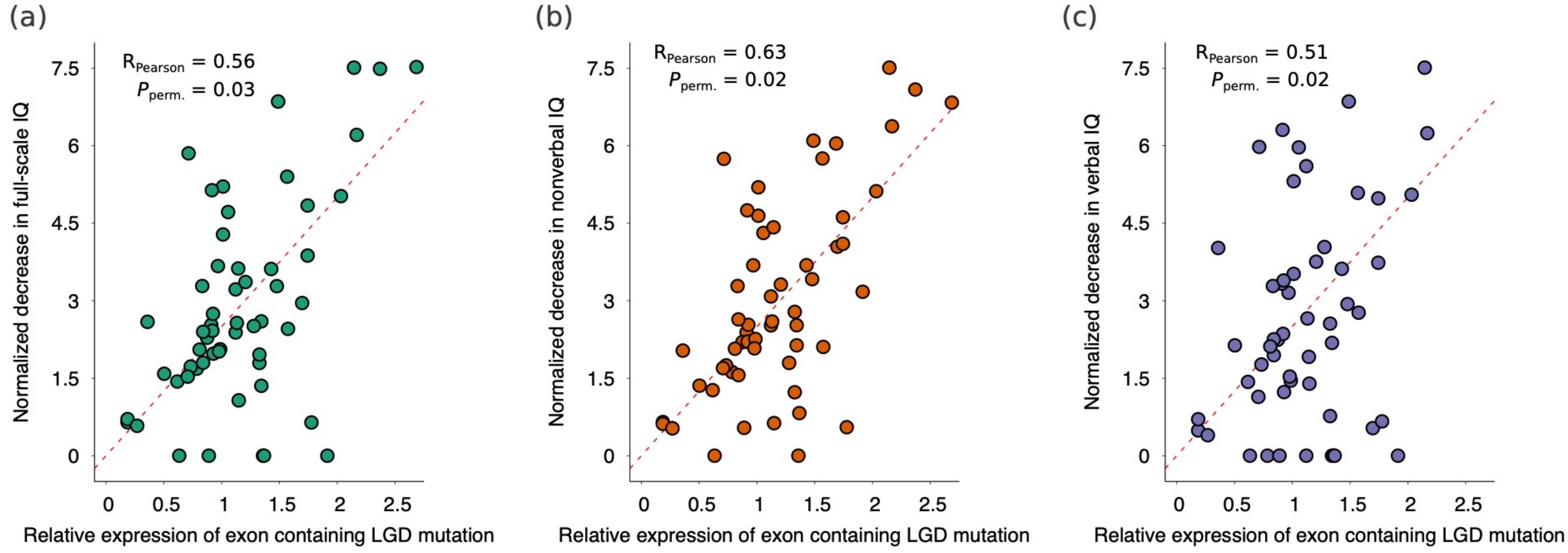


**Supplementary Figure 12:** Relationship between the relative expression of exons harboring LGD mutations and the corresponding decrease in proband's IQ. From left to right, the scatterplots show **(a)** full-scale IQ (FSIQ), **(b)** nonverbal IQ (NVIQ), and **(c)** verbal IQ (VIQ) scores. Each point in the scatterplots corresponds to a proband in SSC affected by an LGD mutation; only genes with recurrent LGD mutations in SSC were considered. The x-axis represents the relative expression, i.e. the ratio of exon expression to total gene expression, of the exon harboring the LGD mutation. The y-axis represents the proband's observed decrease in IQ (relative to wild-type score of 100) normalized by the Phenotype Dosage Sensitivity (PDS) parameter of each gene (see Methods). Red dashed lines represent the linear regression fits across all points.

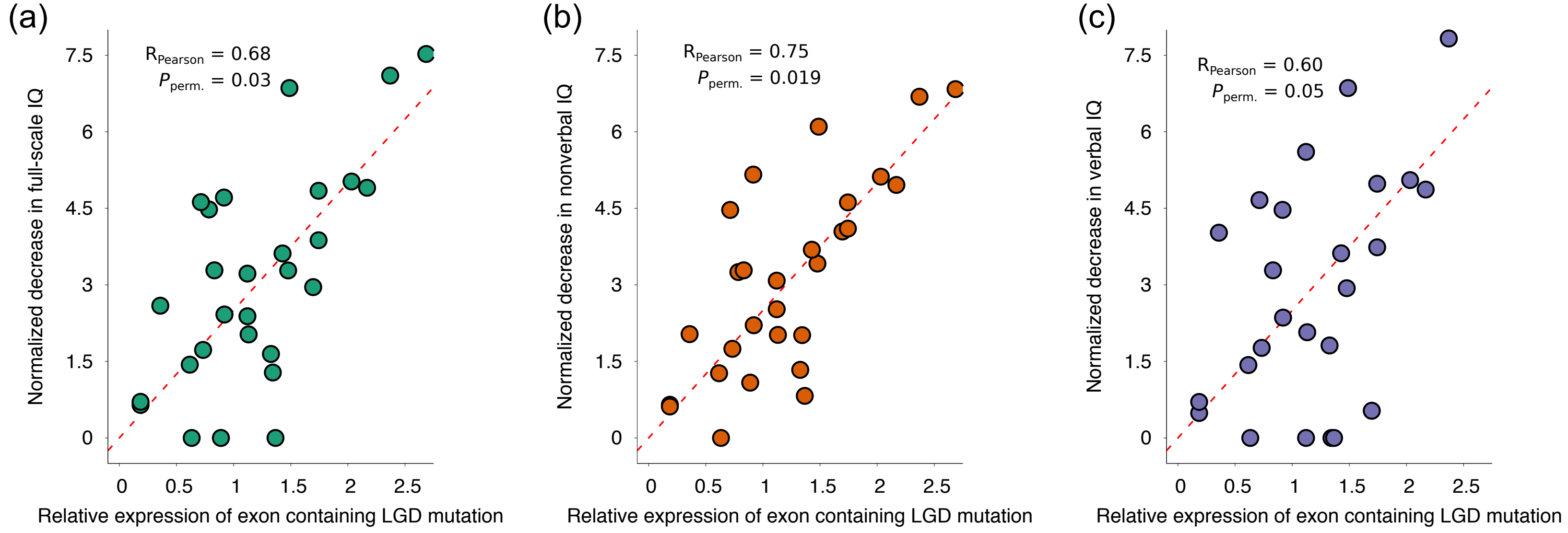


**Supplementary Figure 13:** Relationship between the relative expression of exons harboring LGD mutations and the corresponding decrease in probands' intellectual phenotypes for the older half of probands in SSC (i.e. older than the median age 8.35 years). From left to right, the scatterplots show **(a)** full-scale IQ (FSIQ), **(b)** nonverbal IQ (NVIQ), and **(c)** verbal IQ (VIQ) scores. Each point in the scatterplots corresponds to a proband in SSC affected by an LGD mutation; only genes with recurrent LGD mutations in SSC were considered. The x-axis represents the relative expression, i.e. the ratio of exon expression to total gene expression, of the exon harboring the LGD mutation. The y-axis represents the proband's observed decrease in IQ (relative to wild-type score of 100) normalized by the Phenotype Dosage Sensitivity (PDS) parameter of each gene (see Methods). Red dashed lines represent the linear regression fits across all points.

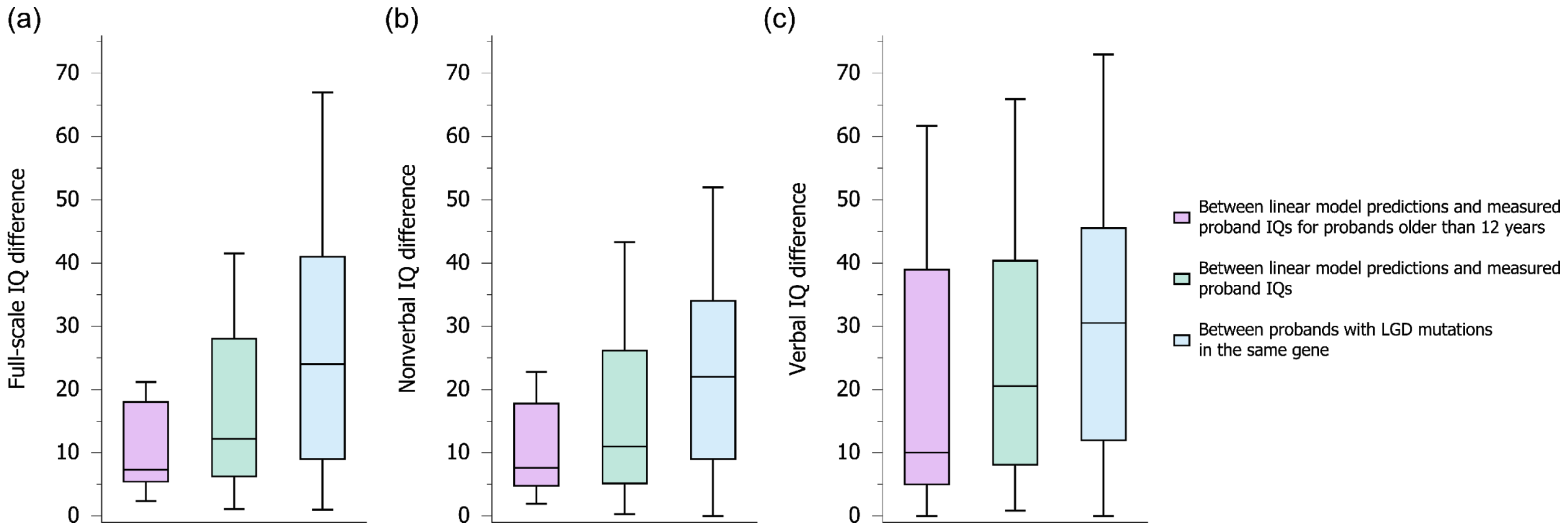


**Supplementary Figure 14:** Distribution of the errors in predicting the effect of LGD mutations on IQ scores based on the linear dosage model. The errors are shown for all probands (green), and for probands older than 12 years (purple); for comparison, the distribution of IQ differences between all probands with LGD mutations in the same genes are also shown (blue). From left to right, plots represent the distribution of score differences for **(a)** full-scale IQ (FSIQ), **(b)** nonverbal IQ (NVIQ), and **(c)** verbal IQ (VIQ). The ends of each solid box represent the upper and lower quartiles, the horizontal lines within each box represent the medians, and the whiskers represent the 5$^{th}$ and 95$^{th}$ percentiles.

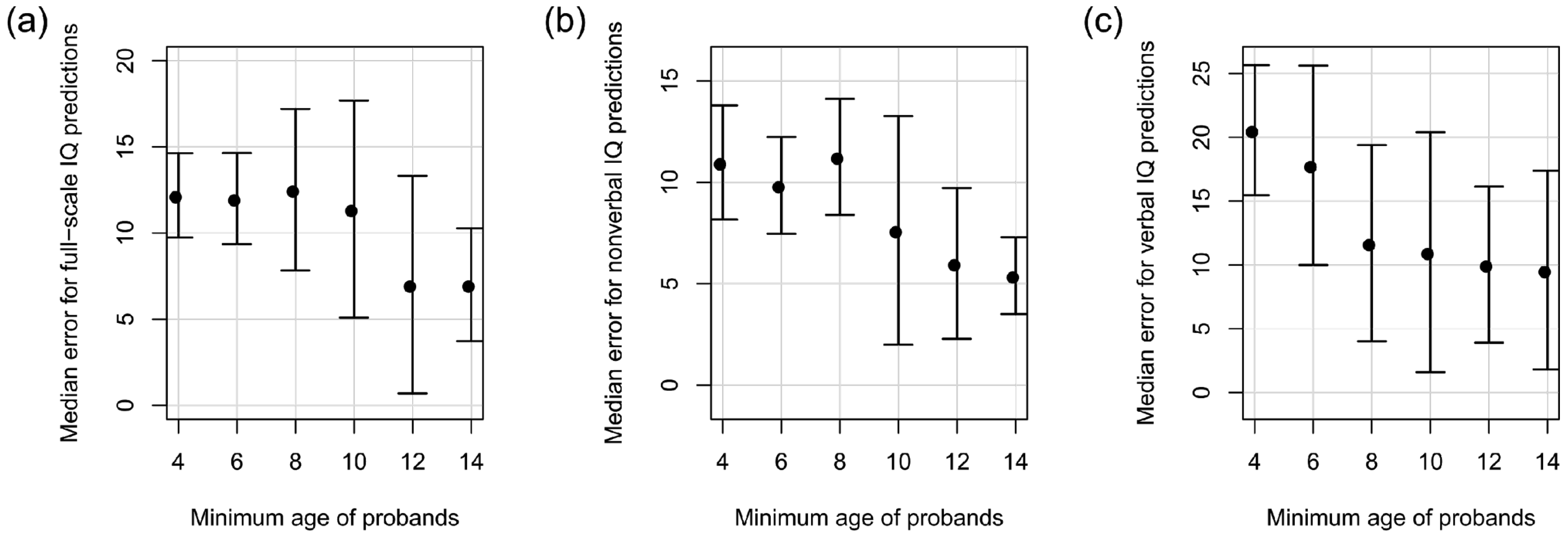


**Supplementary Figure 15:** Median error in predicting the effects of LGD mutations on IQs for older probands in SSC. From left to right, plots represent the median prediction errors for **(a)** full-scale IQ (FSIQ), **(b)** nonverbal IQ (NVIQ), and **(c)** verbal IQ (VIQ). The x-axis represents the minimum age of probands used for leave-one-out predictions. The y-axis represents the median prediction error based on the linear dosage model. Error bars were estimated using bootstrapping.

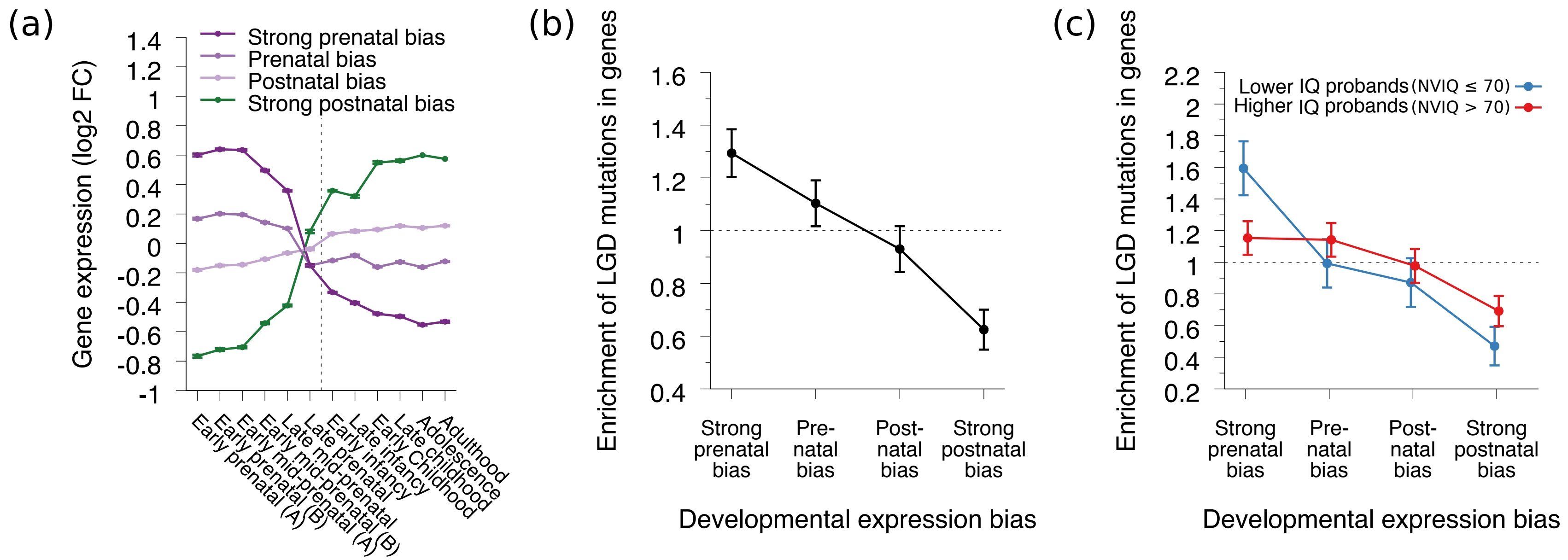


**Supplementary Figure 16**: Relationship between the developmental expression profiles of ASD genes and IQ phenotypes. (**a**) Developmental expression profiles for genes harboring LGD mutations in SSC. Genes were grouped into four bins based on their developmental expression bias, i.e. the fold change between prenatal and postnatal expression. Lines represent the expression for genes in the four bins: genes with strong prenatal bias, genes with prenatal bias, genes with postnatal bias, and genes with strong postnatal bias. The x-axis represents different time periods across human brain development. The y-axis represents the relative expression of genes in each bin, defined as the average log2 fold-change relative to the mean expression level across all periods. Error bars represent the SEM. The vertical grey line delineates prenatal and postnatal developmental periods. (**b**,**c**) Enrichment of LGD mutations in genes across different bins (x-axis). The y-axes represent the enrichment of mutation in each group compared to the expected mutation frequency based on random shuffling of mutations across the full coding length of the corresponding protein. Error bars represent the SEM. (**b**) The enrichment of SSC LGD mutations in genes from the four bins. (**c**) The enrichment of SSC LGD mutations in the four bins for probands with higher (>70, red) and lower (<70, blue) nonverbal IQs.

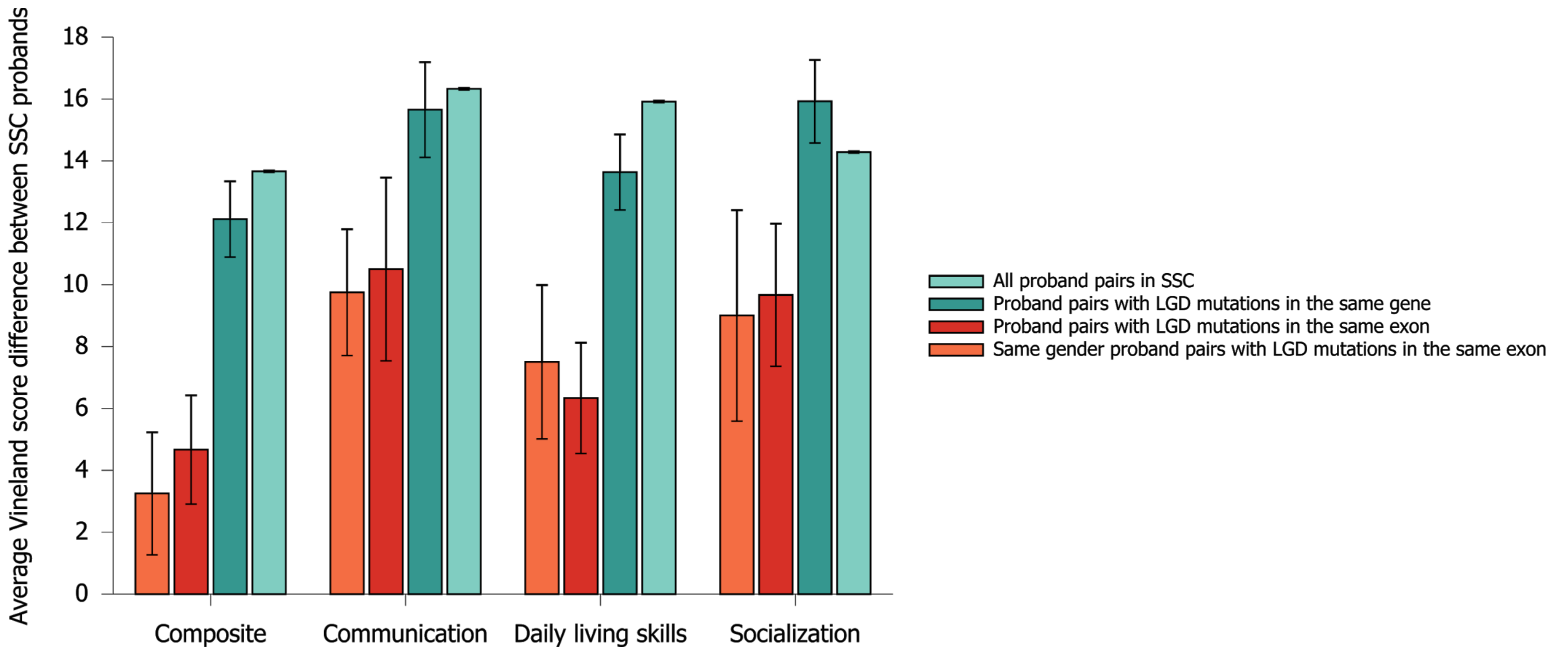


**Supplementary Figure 17:** Average differences in adaptive behavior scores, i.e. Vineland Adaptive Behavior Scales, 2nd ed. (VABS), between SSC probands in SSC. Each bar shows the average difference in VABS scores between pairs of probands. From left to right, bar groups represent differences in the composite score, and in communication, in daily living skills (DLS), and in socialization sub-scores. Within each bar group, bars represent, from right to left, the average score difference between all pairs of probands in the SSC cohort (light green), between probands with *de novo* LGD mutations in the same gene (dark green), between probands with *de novo* LGD mutations in the same exon (red), and between probands of the same gender and with *de novo* LGD mutations in the same exon (orange). Error bars represent the SEM.

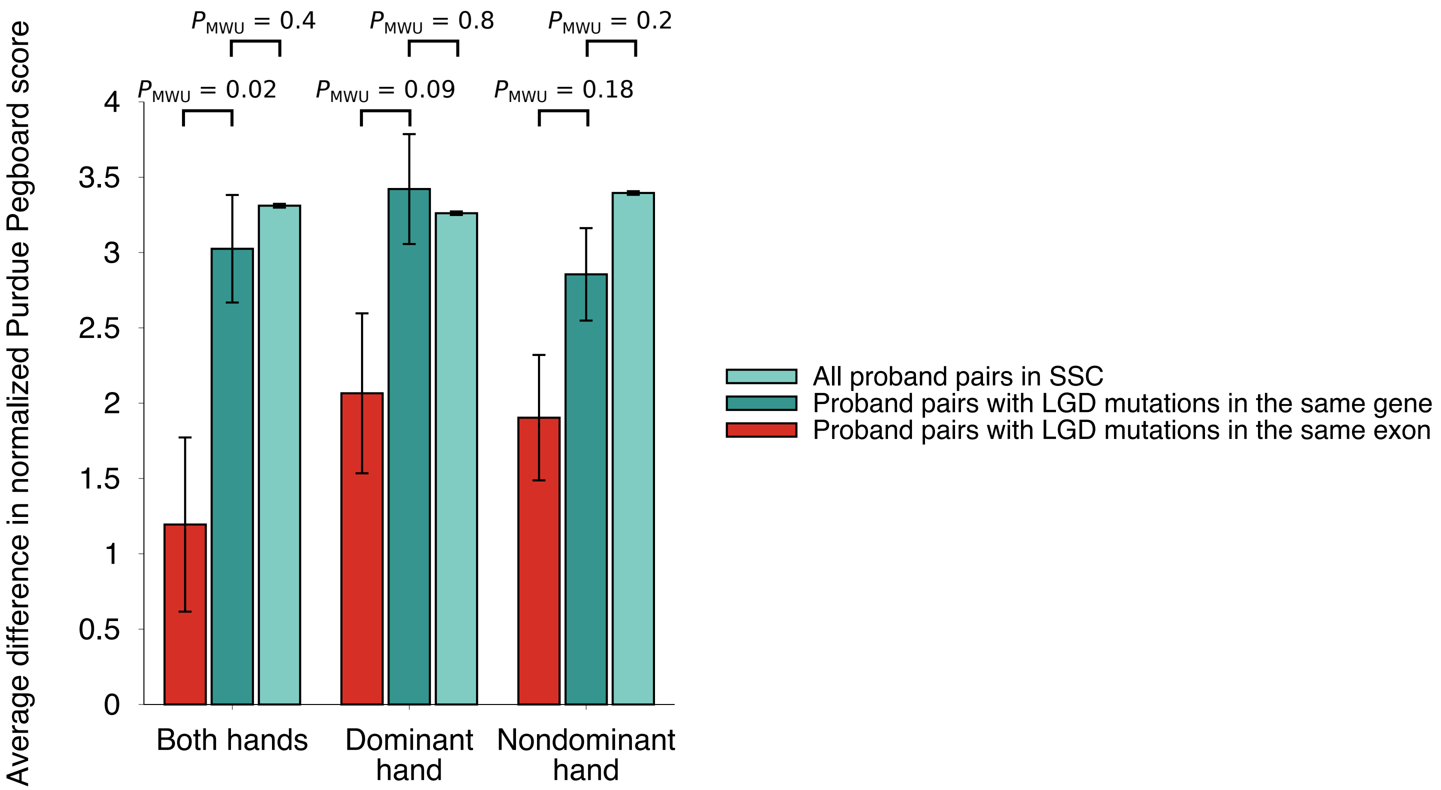


**Supplementary Figure 18:** Average difference in Purdue Pegboard Test scores between pairs of ASD probands. From left to right, bar groups represent Purdue Pegboard Test scores for both hands, for the dominant hand, and for the non-dominant hand. Within each bar group, bars represent, from right to left, the average difference between all pairs of probands in SSC (light green), between pairs of probands with *de novo* LGD mutations in the same gene (dark green), and between pairs of probands with LGD mutations in the same exon (red). Purdue Pegboard scores were adjusted to account for probands' age and gender (see Methods). The statistical significances were calculated using Mann-Whitney U tests ($P_{MWU}$). Error bars represent the SEM.

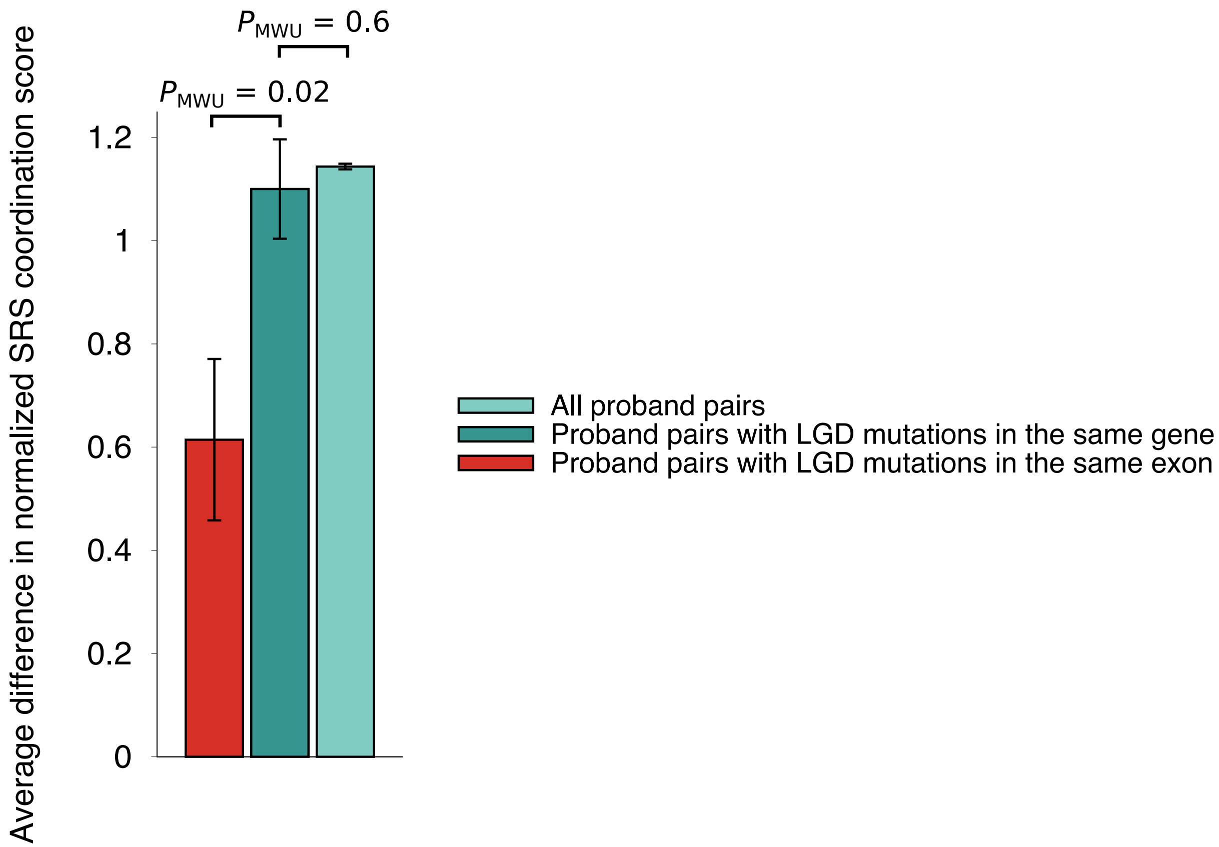


**Supplementary Figure 19:** Average difference in Social Responsiveness Scales (SRS) coordination score between ASD probands. The bars represent, from right to left, all pairs of probands in SSC (light green), pairs of probands with *de novo* LGD mutations in the same gene (dark green), and between pairs of probands with LGD mutations in the same exon (red)**.** SRS scores were adjusted to account for probands' age and gender (see Methods). The statistical significance was determined using Mann-Whitney U tests ($P_{MWU}$). Error bars represent the SEM.

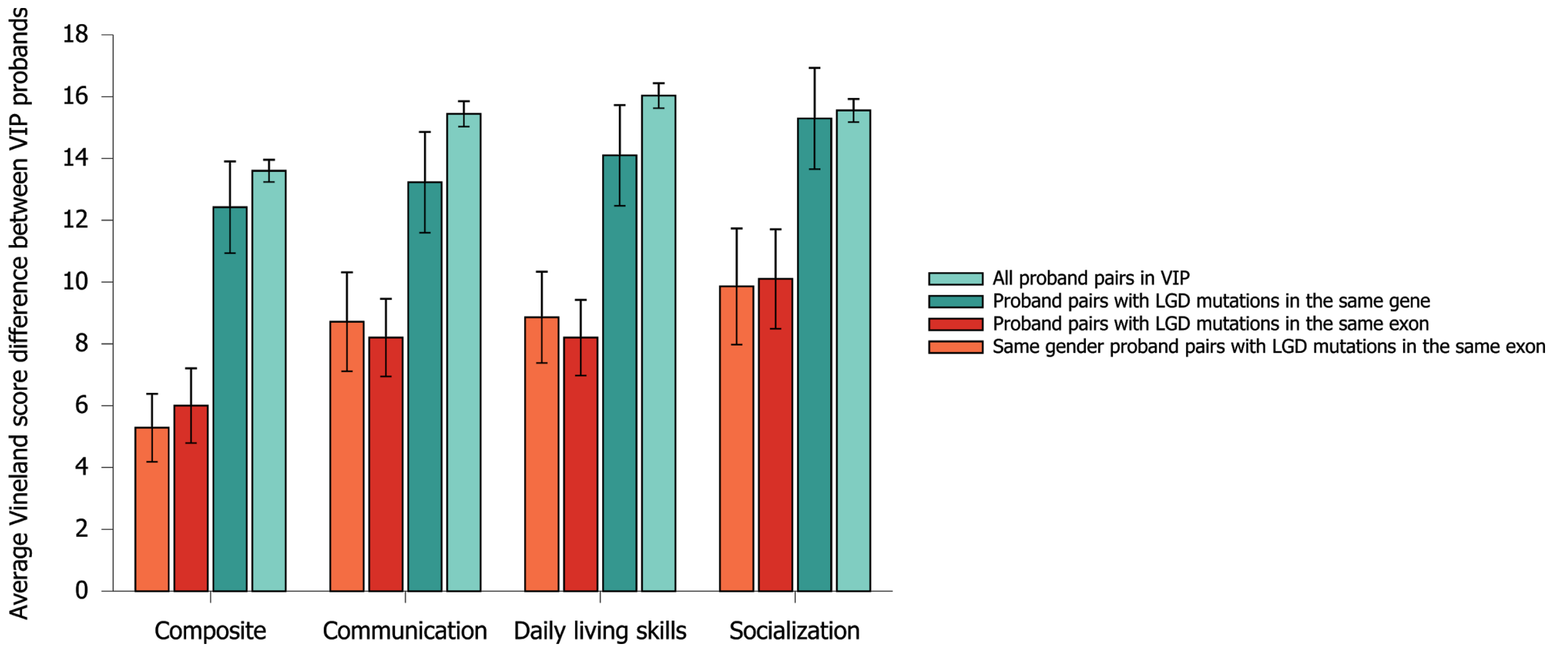


**Supplementary Figure 20:** Average difference in Vineland adaptive behavior scores between probands in the Simons Variation in Individuals Project (VIP). Each bar shows the average difference in Vineland scores between pairs of probands. From left to right, bar groups represent differences in the composite standard score, and in communication, daily living skills (DLS), and socialization subscores. From right to left within each bar group, bars represent the average difference between all pairs of probands in the VIP cohort (light green), between probands with *de novo* LGD mutations in the same gene (dark green), between probands with *de novo* LGD mutations in the same exon (red), and between probands of the same gender with *de novo* LGD mutations in the same exon (orange). Error bars represent the SEM.

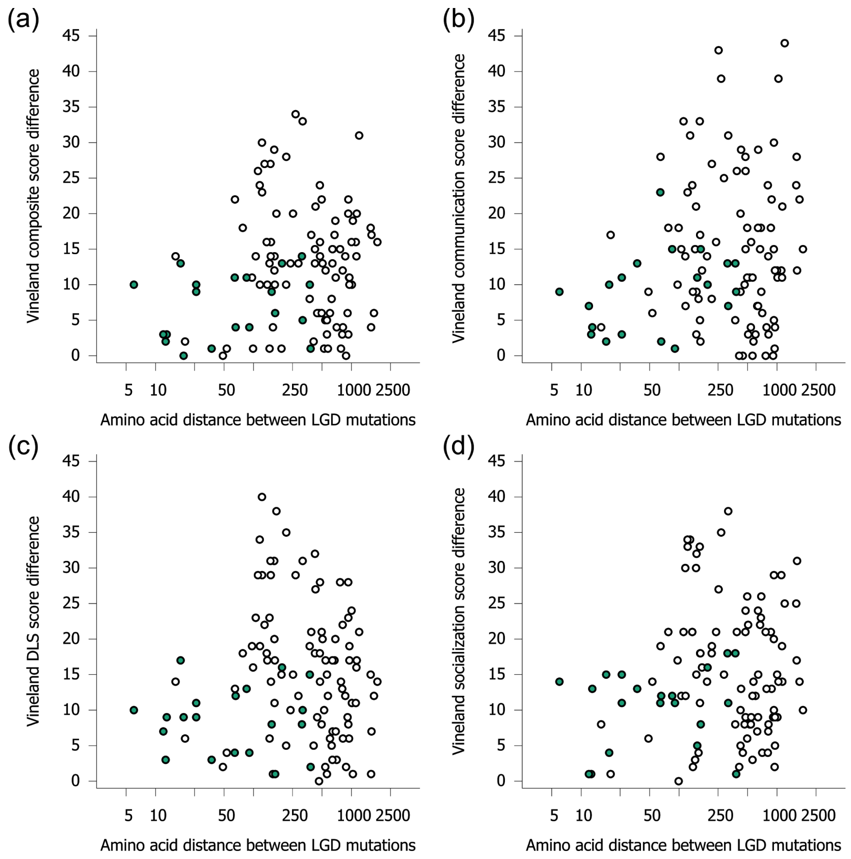


**Supplementary Figure 21:** Amino acid distance between LGD mutations in protein sequence versus the Vineland score differences between probands; data from the SSC and VIP cohorts were combined. Clockwise from top left, plots show differences in **(a)** Vineland composite standard score, **(b)** Vineland communication, **(c)** Vineland daily living skills (DLS), and **(d)** Vineland socialization subscores. In the figures, each point corresponds to a pair of probands affected by *de novo* LGD mutations in the same gene. The x-axis represents the amino acid distance between LGD mutations, and the y-axis represents the difference between the Vineland scores. Open (white) points correspond to proband pairs with mutations in different exons of the same gene; filled (colored) points correspond to pairs with mutations in the same exon.

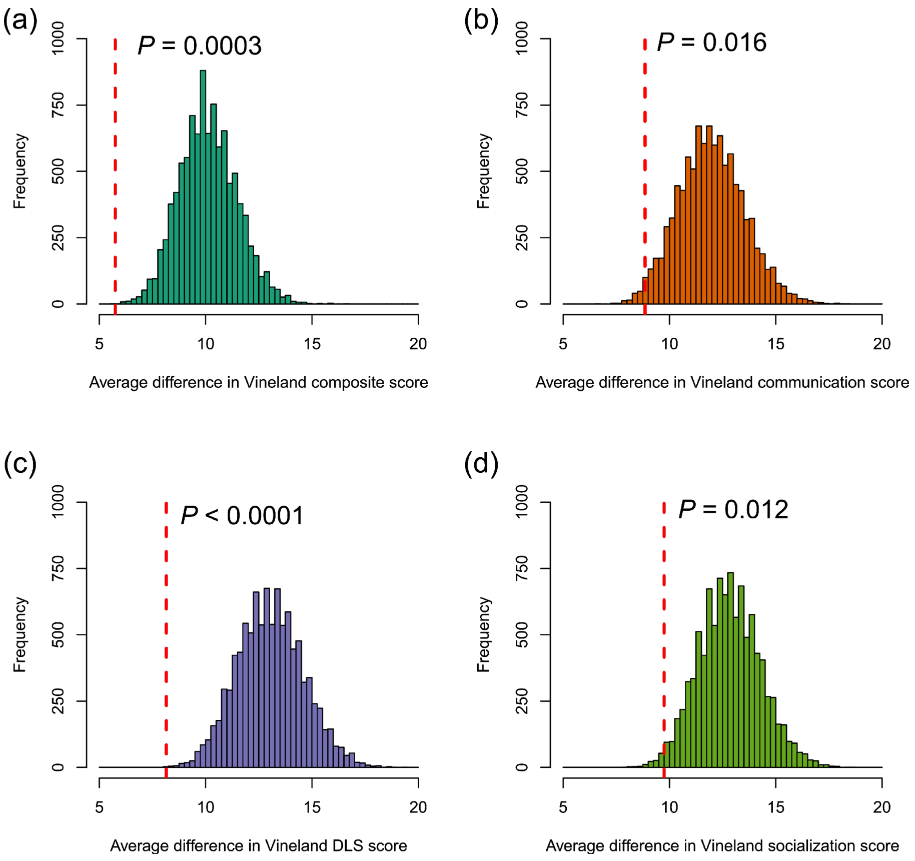


**Supplementary Figure 22:** Simulated null distributions of the average Vineland score difference between ASD probands (from the combined SSC and VIP cohort) with *de novo* LGD mutations in the same gene and separated by similar distances in corresponding protein sequence. Clockwise from top left, histograms represent the distribution of average differences in Vineland for **(a)** the composite standard score, and for **(b)** communication, **(c)** daily living skills (DLS), and **(d)** socialization subscores. Each value in the histogram represents the average Vineland score difference between pairs of probands with mutations in the same gene and with protein amino acid distances similar to the distances empirically observed for LGD mutations in the same exon (see Methods). The red dashed lines represent the observed average score differences between probands with mutations in the same exon.

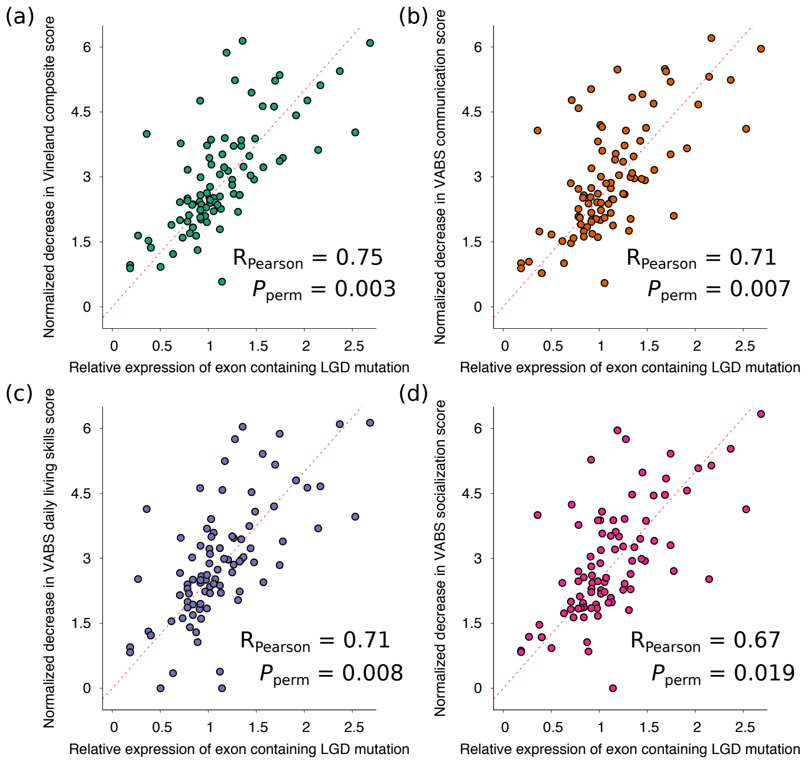


**Supplementary Figure 23:** Relationship between the relative expression of exons harboring LGD mutations and the corresponding decrease in probands' adaptive behavior phenotypes, i.e. Vineland Adaptive Behavior Scales (VABS) in SSC and VIP probands. Clockwise from top left, the scatterplots show standardized VABS **(a)** composite, **(b)** communication, **(c)** daily living skills, and **(d)** socialization scores. Each point in the scatterplots corresponds to a proband in SSC or VIP affected by an LGD mutation; only genes with recurrent LGD mutations across both datasets were considered. The x-axis represents the relative expression, i.e. the ratio of exon expression to total gene expression, of the exon harboring the LGD mutation. The y-axis represents the proband's observed decrease in VABS score (relative to wild-type score of 100) normalized by the Phenotype Dosage Sensitivity (PDS) parameter of each gene (see Methods). Red dashed lines represent the linear regression lines across all points.

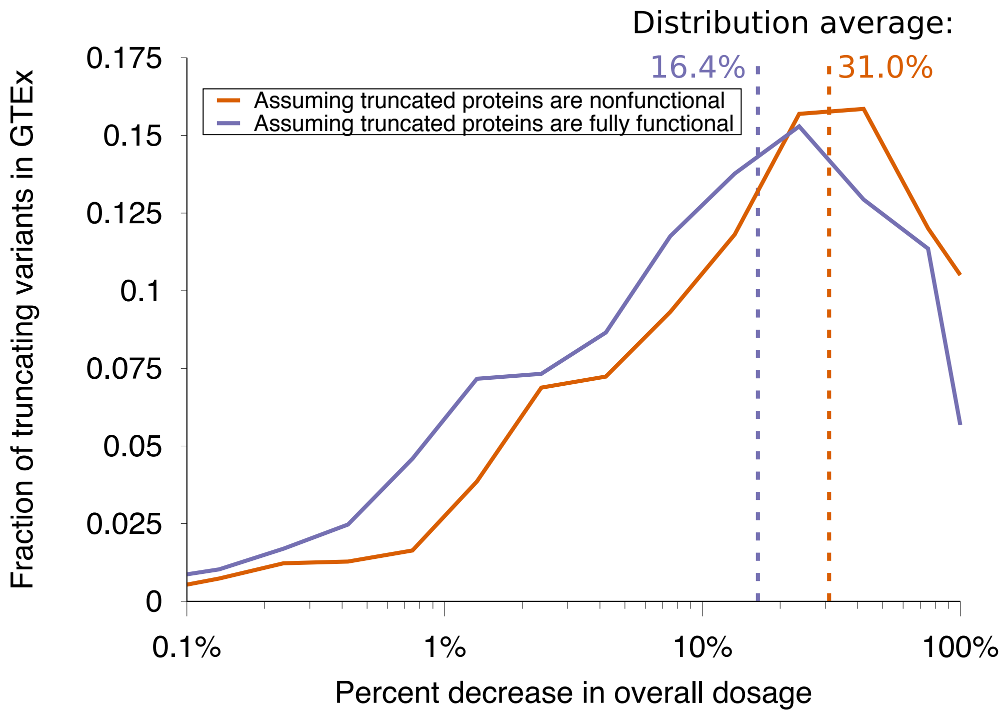


**Supplementary Figure 24:** The distribution of changes in overall gene expression dosage due to LGD variants in GTEx. For each LGD variant in GTEx, we calculated the relative (percent) change in total gene expression caused by the LGD variant in one of 10 major tissues analyzed in GTEx (see Methods); the results were combined across tissues. Each distribution represents the estimated dosage changes across all variants. The distribution shown in purple represents the change in overall gene dosage due to nonsense-mediated decay (NMD), assuming that truncated proteins surviving NMD are fully functional. The distribution in orange represents the change in overall gene dosage assuming that all truncated proteins are nonfunctional. The vertical dashed lines represent the average of each distribution.

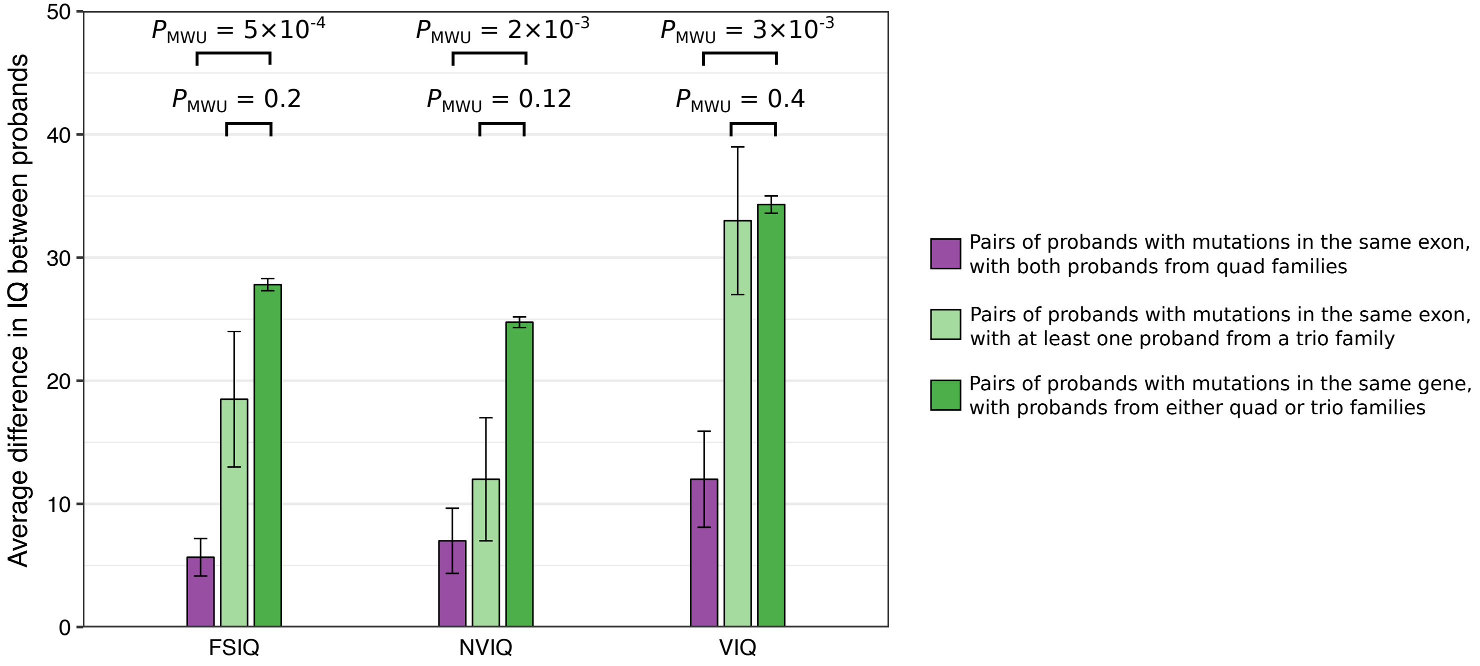


**Supplementary Figure 25:** Average IQ differences between SSC probands from quad and trio ASD families. Quad families have a single affected child among with one or multiple unaffected; trio families have an affected child with no siblings. From left to right, differences are shown for full-scale IQ (FSIQ), nonverbal IQ (NVIQ), and verbal IQ (VIQ) scores. For each score, bars represent the average IQ difference between probands with *de novo* LGD mutations in the same gene (dark green), between pairs of probands with mutations in the same exon and with at least one proband from a trio family (light green), and between probands from quad families with mutations in the same exon (purple). The statistical significance was calculated using Mann-Whitney U test ($P_{MWU}$). Error bars indicate the SEM.

**Supplementary Table 1**: Statistical tests of phenotypic similarity, and associated p-values. Unless indicated otherwise, proband phenotypes were obtained from Simons Simplex Collection (SSC). Only Vineland Adaptive Behavior Scales (VABS) scores were available for both SSC and Simons Variation in Individuals Project (SVIP) cohorts.

| **Phenotypic comparison** | **Statistical test** | **Phenotype** | **P-value; test statistic** |
|---|---|---|---|
| Compare phenotype differences between proband pairs with LGDs in the same gene and between all proband pairs in SSC | Mann-Whitney *U*, one-tail | FSIQ | *P* = 0.2; 11% smaller |
| | | NVIQ | *P* = 0.14; 12% smaller |
| | | VIQ | *P* = 0.5; 1.1% smaller |
| | | VABS | *P* = 0.23; 11% smaller |
| Compare phenotype differences between proband pairs with LGDs ≤ 1 kbp and between proband pairs with LGDs > 1 kbp | Mann-Whitney *U*, one-tail | FSIQ | *P* = 0.002 |
| | | NVIQ | *P* = 0.005 |
| | | VIQ | *P* = 0.01 |
| Correlate phenotype differences and LGD mutation proximity | Spearman correlation | FSIQ | *P* = 0.5; ρ = 0.09 |
| | | NVIQ | *P* = 0.4; ρ = 0.1 |
| | | VIQ | *P* = 0.8; ρ = 0.03 |
| Compare phenotype differences between proband pairs with LGDs in the same exon and between proband pairs with LGDs in different exons of the same gene | Mann-Whitney *U*, one-tail | FSIQ | *P* = 0.003 |
| | | NVIQ | *P* = 0.005 |
| | | VIQ | *P* = 0.016 |
| | | SSC VABS | *P* = 0.017 |
| | | Purdue Pegboard | *P* = 0.02 |
| | | SRS | *P* = 0.05 |
| | | SVIP VABS | *P* = 0.014 |
| Compare phenotype differences between proband pairs with LGDs in neighboring | Mann-Whitney *U*, one-tail | FSIQ | *P* = 0.6 |

| | | | |
|---|---|---|---|
| exons and between proband pairs with LGDs in non-neighboring exons | | NVIQ | *P* = 0.18 |
| | | VIQ | *P* = 0.8 |
| | | SSC VABS | *P* = 0.14 |
| | | SVIP VABS | *P* = 0.6 |
| Compare phenotype differences between pairs of probands of the same gender with LGDs in the same exon and between pairs of probands of different gender with LGDs in the same exon | Mann-Whitney *U*, one-tail | FSIQ | *P* = 0.04 |
| | | NVIQ | *P* = 0.29 |
| | | VIQ | *P* = 0.07 |
| Compare phenotype differences between proband pairs with LGDs in the same exon and between proband pairs with LGDs in different exons with similar amino acid distances | Permutation test, conditioned on similar distance between mutations | FSIQ | *P* = 0.01 |
| | | NVIQ | *P* = 0.002 |
| | | VIQ | *P* = 0.018 |
| | | SSC/SVIP VABS | *P* = 3e-4 |
| Correlate fraction of protein truncated by LGD and proband phenotype | Pearson correlation | FSIQ | *P* = 0.35; R = 0.05 |
| | | NVIQ | *P* = 0.35; R = 0.05 |
| | | VIQ | *P* = 0.28; R = 0.06 |
| | | SSC VABS | *P* = 0.47; R = -0.04 |
| | | SVIP VABS | *P* = 0.7; R = -0.08 |
| Compare phenotypes of probands with a smaller fraction of truncated sequence and with a larger fraction of truncated sequence for the same protein | Wilcoxon signed-rank, one-tail | FSIQ | *P* = 0.9 |
| | | NVIQ | *P* = 0.44 |
| | | VIQ | *P* = 0.89 |
| Compare phenotype differences between probands pairs with LGDs in the same exon and same protein domain and between probands pairs with LGDs in the same exon but in different domains of the same protein | Mann-Whitney *U*, one-tail | FSIQ | *P* = 0.9 |
| | | NVIQ | *P* = 0.7 |
| | | VIQ | *P* = 0.8 |

| | | | |
|---|---|---|---|
| Compare observed expression-phenotype correlation among LGDs and expected correlations based on permuted exon data | Permutation test | FSIQ | *P* = 0.03; R = 0.56 |
| | | NVIQ | *P* = 0.02; R = 0.63 |
| | | VIQ | *P* = 0.02; R = 0.51 |
| | | FSIQ, age > 8 yr | *P* = 0.03; R = 0.68 |
| | | NVIQ, age > 8 yr | *P* = 0.019; R = 0.75 |
| | | VIQ, age > 8 yr | *P* = 0.05; R = 0.60 |
| Compare phenotypic prediction errors for the linear dosage model and for predictions based on the assumption that LGD mutations in the same gene lead to the same phenotypic consequences | Mann-Whitney *U*, one-tail | FSIQ | *P* = 0.019; error = 12.2 |
| | | NVIQ | *P* = 0.014; error = 11.0 |
| | | VIQ | *P* = 0.017; error = 20.6 |
| | | FSIQ, age > 12 yr | error = 7.0 |
| | | NVIQ, age > 12 yr | error = 7.6 |
| | | VIQ, age > 12 yr | error = 10.0 |
| Compare phenotypic prediction errors for the linear dosage model, for predictions based on probands of the same gender versus predictions based on probands of different genders | Mann-Whitney *U*, one-tail | FSIQ | *P* = 0.003; error = 11.1 |
| | | NVIQ | *P* = 0.018; error = 9.1 |
| | | VIQ | *P* = 0.02; error = 15.9 |

**Supplementary Table 2**: Predicted loss of gene expression dosage for each LGD mutation.

| **Proband** | **Variant** | **Gene** | **Predicted Loss of expression dosage (fraction)** |
|---|---|---|---|
| 13585.p1 | 12:109577549:A:AG | ACACB | 0.10826519 |
| 13702.p1 | 3:58491004:CA:C | ACOX2 | 0.20175851 |
| 12561.p1 | 20:3655475:G:A | ADAM33 | 0.13312044 |
| 14490.p1 | 3:64547409:G:A | ADAMTS9 | 0.21079264 |
| 11717.p1 | 16:4165336:CT:C | ADCY9 | 0.13125763 |
| 12130.p1 | 20:49510027:CTT:C | ADNP | 0.16505754 |
| 13545.p1 | 20:49509094:G:GT | ADNP | 0.16505754 |
| 12246.p1 | 4:74357772:C:T | AFM | 0.14623134 |
| 13695.p1 | 1:27876868:G:GT | AHDC1 | 0.15756625 |
| 14474.p1 | 7:91726153:C:T | AKAP9 | 0.30853464 |
| 13993.p1 | 9:116824955:A:G | AMBP | 0.16066873 |
| 11277.p1 | 11:94599334:G:T | AMOTL1 | 0.11136076 |
| 12645.p1 | 4:114232545:C:T | ANK2 | 0.1513039 |
| 13768.p1 | 4:114277599:AG:A | ANK2 | 0.13569239 |
| 14256.p1 | 4:114251469:C:T | ANK2 | 0.1414001 |
| 12854.p1 | 17:54558136:CAG:C | ANKFN1 | 0.16808306 |
| 12507.p1 | 16:89350771:CTTTG:C | ANKRD11 | 0.15543117 |
| 13905.p1 | 16:89351042:GTGTTT:G | ANKRD11 | 0.15543117 |
| 12792.p1 | 11:22276997:C:T | ANO5 | 0.17211645 |
| 14114.p1 | 11:428173:C:CT | ANO9 | 0.14948482 |
| 14535.p1 | 11:6417424:G:GC | APBB1 | 0.1674499 |
| 11466.p1 | 1:161017762:G:A | ARHGAP30 | 0.19631841 |
| 13097.p1 | 14:32561337:G:T | ARHGAP5 | 0.15851713 |
| 13447.p1 | 6:157527664:CTGTT:C | ARID1B | 0.30868295 |
| 14393.p1 | 6:157510805:A:AC | ARID1B | 0.19893591 |
| 14405.p1 | 22:19965564:G:A | ARVCF | 0.20282514 |

| | | | |
|---|---|---|---|
| 11715.p1 | 10:52005130:T:TG | ASAH2 | 0.10510733 |
| 13678.p1 | 1:155340680:C:A | ASH1L | 0.19324424 |
| 13616.p1 | 4:47571000:T:TG | ATP10D | 0.15164379 |
| 14003.p1 | 1:116916864:G:A | ATP1A1 | 0.10147881 |
| 12840.p1 | 1:169096506:C:T | ATP1B1 | 0.19699168 |
| 14070.p1 | 12:58025102:G:GC | B4GALNT1 | 0.15654 |
| 13634.p1 | 8:143605599:C:G | BAI1 | 0.20572809 |
| 14581.p1 | 2:160182250:G:GT | BAZ2B | 0.30256011 |
| 13183.p1 | 2:60689253:A:AG | BCL11A | 0.14563672 |
| 14210.p1 | 5:172571588:C:T | BNIP1 | 0.11627822 |
| 12610.p1 | 17:41197776:C:T | BRCA1 | 0.17735247 |
| 13618.p1 | 8:37702145:CAG:C | BRF2 | 0.15455685 |
| 14292.p1 | 21:40568453:GT:G | BRWD1 | 0.11455266 |
| 14012.p1 | 19:17516098:A:G | BST2 | 0.17586677 |
| 11025.p1 | 10:93723898:CA:C | BTAF1 | 0.13791472 |
| 14442.p1 | 6:38548004:G:A | BTBD9 | 0.15001187 |
| 11132.p1 | 6:26508728:G:A | BTN1A1 | 0.15833001 |
| 14563.p1 | 10:128193561:A:AG | C10orf90 | 0.19447859 |
| 11256.p1 | 11:68029416:AGTCTCTACCT:A | C11orf24 | 0.20290453 |
| 11208.p1 | 16:685776:T:G | C16orf13 | 0.13161974 |
| 14531.p1 | 21:45750792:G:A | C21orf2 | 0.15095461 |
| 13526.p1 | 3:54921984:A:G | CACNA2D3 | 0.15100684 |
| 13977.p1 | 1:223853218:AC:A | CAPN8 | 0.1029181 |
| 12161.p1 | 13:111290774:C:T | CARKD | 0.18434948 |
| 14631.p1 | 6:90581006:A:G | CASP8AP2 | 0.18421756 |
| 13234.p1 | 17:77809050:G:A | CBX4 | 0.15315664 |
| 12854.p1 | 19:14031646:G:T | CC2D1A | 0.1487455 |
| 14575.p1 | 9:15784693:G:A | CCDC171 | 0.41979739 |
| 13060.p1 | 3:56627767:AAAGT:A | CCDC66 | 0.24117485 |
| 14203.p1 | 11:66358293:C:A | CCDC87 | 0.14788699 |

| | | | |
|---|---|---|---|
| 12622.p1 | 15:43020870:CT:C | CDAN1 | 0.15170339 |
| 13809.p1 | 5:137534408:CTG:C | CDC23 | 0.14847739 |
| 13606.p1 | 14:103434646:G:A | CDC42BPB | 0.14683506 |
| 14559.p1 | 5:98194693:C:CA | CHD1 | 0.22913829 |
| 13614.p1 | 15:93563244:C:T | CHD2 | 0.10829542 |
| 13618.p1 | 15:93524060:TAAAG:T | CHD2 | 0.13294246 |
| 13818.p1 | 15:93563282:T:TG | CHD2 | 0.10829542 |
| 11654.p1 | 14:21871373:T:C | CHD8 | 0.14860098 |
| 12752.p1 | 14:21861376:ACT:A | CHD8 | 0.20617996 |
| 12991.p1 | 14:21861643:TCTTC:T | CHD8 | 0.15608253 |
| 13844.p1 | 14:21871178:G:A | CHD8 | 0.14860098 |
| 13900.p1 | 14:21895989:ACTCTTGCACGTCCCATCACAGTAGCAAGGAGTACTCACTTGAGCTTG:A | CHD8 | 0.13194235 |
| 14016.p1 | 14:21870169:G:A | CHD8 | 0.15714353 |
| 14233.p1 | 14:21859175:A:AT | CHD8 | 0.24581399 |
| 12642.p1 | 19:42792016:C:T | CIC | 0.15477201 |
| 13254.p1 | 2:6990025:G:A | CMPK2 | 0.21029353 |
| 12444.p1 | 19:54656675:CTG:C | CNOT3 | 0.28668018 |
| 12118.p1 | 4:109790273:GT:G | COL25A1 | 0.14780087 |
| 12219.p1 | 19:10091481:C:T | COL5A3 | 0.17163782 |
| 11685.p1 | 3:130345392:G:A | COL6A6 | 0.14275299 |
| 12066.p1 | 17:28770924:C:G | CPD | 0.15307182 |
| 13608.p1 | 1:115280664:G:A | CSDE1 | 0.1808028 |
| 12030.p1 | 1:34631390:CG:C | CSMD2 | 0.10172333 |
| 13439.p1 | 10:53458249:T:TA | CSTF2T | 0.14788699 |
| 14346.p1 | 16:67650748:C:CA | CTCF | 0.15529216 |
| 13829.p1 | 10:68535201:C:A | CTNNA3 | 0.17291486 |
| 12211.p1 | 3:41275346:G:A | CTNNB1 | 0.27455511 |
| 13070.p1 | 7:117423007:TCA:T | CTTNBP2 | 0.15625663 |
| 13844.p1 | 10:17127755:G:A | CUBN | 0.12287071 |
| 11452.p1 | 2:225376218:C:A | CUL3 | 0.29199636 |

| | | | |
|---|---|---|---|
| 14307.p1 | 2:127944869:AC:A | CYP27C1 | 0.15330893 |
| 11999.p1 | X:41204655:A:G | DDX3X | 0.16869507 |
| 11028.p1 | 4:169138089:CTTTG:C | DDX60 | 0.19138275 |
| 14578.p1 | 5:54563046:G:A | DHX29 | 0.21997131 |
| 13012.p1 | 21:47958429:A:ACTGGTCT | DIP2A | 0.19794856 |
| 13106.p1 | 21:47957426:G:A | DIP2A | 0.21927654 |
| 12705.p1 | 10:428608:G:GC | DIP2C | 0.14594121 |
| 12653.p1 | 6:170593075:C:CA | DLL1 | 0.15654953 |
| 11610.p1 | 5:13923403:CA:C | DNAH5 | 0.21325696 |
| 12311.p1 | 2:25505356:A:AT | DNMT3A | 0.11085025 |
| 14613.p1 | 8:25257428:C:CT | DOCK5 | 0.15268659 |
| 12093.p1 | 19:2214494:C:T | DOT1L | 0.12501116 |
| 13630.p1 | 2:32168370:C:T | DPY30 | 0.10239607 |
| 12329.p1 | 21:41457527:A:T | DSCAM | 0.17064055 |
| 13735.p1 | 21:41414330:A:AT | DSCAM | 0.26909874 |
| 14597.p1 | 21:41457640:T:TTA | DSCAM | 0.17064055 |
| 13582.p1 | 11:117301436:A:G | DSCAML1 | 0.2326459 |
| 11042.p1 | 18:29055782:T:TA | DSG3 | 0.16051888 |
| 13612.p1 | 6:56393643:GGTTT:G | DST | 0.25249311 |
| 11242.p1 | 3:183887912:G:GT | DVL3 | 0.1584368 |
| 12099.p1 | 21:38845116:CAT:C | DYRK1A | 0.14596027 |
| 13256.p1 | 21:38877891:AGGTCTGTGCTGCTGC:A | DYRK1A | 0.1694401 |
| 13552.p1 | 21:38877833:GC:G | DYRK1A | 0.1694401 |
| 13890.p1 | 21:38865466:G:A | DYRK1A | 0.14293675 |
| 14308.p1 | 1:21584017:C:T | ECE1 | 0.15513179 |
| 13364.p1 | 9:95277357:G:A | ECM2 | 0.16543499 |
| 13590.p1 | 17:28400678:GT:G | EFCAB5 | 0.13516775 |
| 14085.p1 | 2:37364142:G:A | EIF2AK2 | 0.10540572 |
| 14248.p1 | 9:23692707:CACGGATG:C | ELAVL2 | 0.16676747 |
| 13128.p1 | 1:23236941:C:T | EPHB2 | 0.19279675 |

| | | | |
|---|---|---|---|
| 12752.p1 | 19:51848515:C:A | ETFB | 0.16535091 |
| 14537.p1 | 2:131810589:C:G | FAM168B | 0.12675843 |
| 12429.p1 | 4:15689874:G:A | FAM200B | 0.15246122 |
| 12685.p1 | 6:17606162:C:T | FAM8A1 | 0.12563541 |
| 12221.p1 | 8:124787443:C:T | FAM91A1 | 0.1723464 |
| 12235.p1 | 13:99100552:AAG:A | FARP1 | 0.18049883 |
| 13810.p1 | 2:48035295:CAG:C | FBXO11 | 0.22887161 |
| 11318.p1 | 10:5979124:C:T | FBXO18 | 0.21153595 |
| 13895.p1 | 5:171326970:G:A | FBXW11 | 0.14874203 |
| 11524.p1 | 1:159778750:G:A | FCRL6 | 0.12700464 |
| 13471.p1 | 1:152286919:CCT:C | FLG | 0.14871216 |
| 11403.p1 | 3:71026115:G:A | FOXP1 | 0.33517698 |
| 12817.p1 | 3:71050170:C:CT | FOXP1 | 0.21016772 |
| 11074.p1 | 4:144620325:G:A | FREM3 | 0.13621475 |
| 13548.p1 | 11:11314689:TG:T | GALNT18 | 0.23737577 |
| 11232.p1 | 7:100283002:TCCCCATCCCCGTTTGTC:T | GIGYF1 | 0.15686257 |
| 14530.p1 | 7:100282216:C:CCATCT | GIGYF1 | 0.16596216 |
| 14533.p1 | 2:233675982:C:T | GIGYF2 | 0.10983416 |
| 13096.p1 | 7:150164231:GA:G | GIMAP8 | 0.13715442 |
| 14287.p1 | 19:48255802:GCACC:G | GLTSCR2 | 0.14676309 |
| 13047.p1 | 14:93286182:G:C | GOLGA5 | 0.19653619 |
| 13551.p1 | 6:117900140:G:A | GOPC | 0.13766054 |
| 11518.p1 | 5:176026121:TCAAAGACCCAGGA:T | GPRIN1 | 0.14965863 |
| 13504.p1 | 15:72461298:C:CT | GRAMD2 | 0.1079721 |
| 11691.p1 | 12:14019043:T:TG | GRIN2B | 0.14261224 |
| 12547.p1 | 12:13764762:C:T | GRIN2B | 0.14366903 |
| 12681.p1 | 12:13722953:T:C | GRIN2B | 0.14057968 |
| 13711.p1 | 14:31613330:G:GT | HECTD1 | 0.12937639 |
| 11940.p1 | 1:42048715:C:CG | HIVEP3 | 0.14018519 |
| 14604.p1 | 2:177054587:A:AT | HOXD1 | 0.17958789 |

| | | | |
|---|---|---|---|
| 11909.p1 | 19:18288571:C:G | IFI30 | 0.15464243 |
| 14441.p1 | 1:79093807:T:A | IFI44L | 0.10739038 |
| 13771.p1 | 9:21207037:A:T | IFNA10 | 0.14788699 |
| 14226.p1 | 13:21205235:T:G | IFT88 | 0.10816319 |
| 13077.p1 | 1:117142752:G:A | IGSF3 | 0.14291455 |
| 13256.p1 | 22:17590329:TC:T | IL17RA | 0.15201509 |
| 13647.p1 | 5:131398057:GT:G | IL3 | 0.13631637 |
| 14621.p1 | 1:153637720:G:A | ILF2 | 0.20145283 |
| 14687.p1 | 13:51948834:G:A | INTS6 | 0.54891003 |
| 12752.p1 | 5:75902128:C:T | IQGAP2 | 0.14582661 |
| 13382.p1 | 14:77492033:GT:G | IRF2BPL | 0.14788699 |
| 11592.p1 | 3:20161089:G:A | KAT2B | 0.15550853 |
| 12108.p1 | 8:41906209:TTTTG:T | KAT6A | 0.13715969 |
| 11008.p1 | 18:44589733:G:A | KATNAL2 | 0.17172417 |
| 11872.p1 | 18:44603833:G:C | KATNAL2 | 0.24328273 |
| 12418.p1 | 1:202705466:G:A | KDM5B | 0.17232576 |
| 13664.p1 | 1:202698172:G:A | KDM5B | 0.18159799 |
| 11329.p1 | 17:7749188:A:G | KDM6B | 0.12790153 |
| 12683.p1 | 17:7749921:CG:C | KDM6B | 0.13823415 |
| 13346.p1 | 17:26961045:G:A | KIAA0100 | 0.14263227 |
| 12653.p1 | 4:6843914:C:T | KIAA0232 | 0.12181146 |
| 14069.p1 | 11:93463107:C:CA | KIAA1731 | 0.43034725 |
| 12728.p1 | 12:39763593:G:GT | KIF21A | 0.13379753 |
| 13849.p1 | 4:39409045:T:TC | KLB | 0.10303443 |
| 11838.p1 | 7:129767258:C:T | KLHDC10 | 0.13986069 |
| 11145.p1 | 11:118344386:TG:T | KMT2A | 0.12250806 |
| 11148.p1 | 7:151842339:G:T | KMT2C | 0.29422251 |
| 12952.p1 | 7:104748100:TC:T | KMT2E | 0.18075503 |
| 14299.p1 | 7:104702706:AC:A | KMT2E | 0.1153122 |
| 13739.p1 | X:153135038:TCA:T | L1CAM | 0.17973276 |

| | | | |
|---|---|---|---|
| 14093.p1 | 10:876865:TCA:T | LARP4B | 0.13188068 |
| 14581.p1 | 17:33313013:CT:C | LIG3 | 0.21576495 |
| 13092.p1 | 19:49004780:C:CAGGTCAG | LMTK3 | 0.21299823 |
| 14477.p1 | 6:40400210:CA:C | LRFN2 | 0.1548691 |
| 12409.p1 | 2:170042308:G:A | LRP2 | 0.13061994 |
| 11121.p1 | 12:12311817:CA:C | LRP6 | 0.14424431 |
| 14587.p1 | 2:238617256:G:GA | LRRFIP1 | 0.18836257 |
| 11843.p1 | 21:30318550:C:A | LTN1 | 0.14875986 |
| 12346.p1 | 2:149225965:GTC:G | MBD5 | 0.18194216 |
| 13621.p1 | 6:90489930:C:T | MDN1 | 0.11151023 |
| 13813.p1 | 17:60108956:AG:A | MED13 | 0.13825349 |
| 12969.p1 | 12:116418553:A:AC | MED13L | 0.12706607 |
| 14416.p1 | 12:116424952:C:T | MED13L | 0.176944 |
| 14075.p1 | 1:3519049:AC:A | MEGF6 | 0.11606531 |
| 13168.p1 | 11:119214624:CT:C | MFRP | 0.11182957 |
| 11776.p1 | 8:145735146:A:AAGCTGGGGGCCGCGCT | MFSD3 | 0.1334405 |
| 13289.p1 | 9:5892550:G:T | MLANA | 0.24717884 |
| 14310.p1 | 21:37744772:CAG:C | MORC3 | 0.27751288 |
| 12473.p1 | 1:113241092:CG:C | MOV10 | 0.14212682 |
| 12224.p1 | 13:20235898:C:T | MPHOSPH8 | 0.14609696 |
| 11986.p1 | 7:24720140:GT:G | MPP6 | 0.23246093 |
| 12948.p1 | 3:135871025:TCAGA:T | MSL2 | 0.15797408 |
| 11521.p1 | 8:17612610:G:C | MTUS1 | 0.10656691 |
| 13896.p1 | 6:30954871:ACGACCTCCAGTGGGGC:A | MUC21 | 0.19451838 |
| 11078.p1 | 11:1253773:C:A | MUC5B | 0.12313592 |
| 13000.p1 | 17:8424204:GA:G | MYH10 | 0.14133837 |
| 14068.p1 | 17:10432099:G:A | MYH2 | 0.12808126 |
| 11577.p1 | 15:59430484:G:A | MYO1E | 0.20314307 |
| 13060.p1 | 19:13246969:T:C | NACC1 | 0.13697495 |
| 11356.p1 | 8:144659231:C:T | NAPRT1 | 0.14743868 |

| | | | |
|---|---|---|---|
| 12805.p1 | 15:60768343:AG:A | NARG2 | 0.24455966 |
| 13761.p1 | 13:36129146:C:T | NBEA | 0.15637915 |
| 12764.p1 | 2:183791570:C:A | NCKAP1 | 0.11610149 |
| 14030.p1 | 2:183866861:C:CCA | NCKAP1 | 0.13866029 |
| 11209.p1 | 4:115760701:G:A | NDST4 | 0.16092553 |
| 14122.p1 | 6:11192635:TGAAAACA:T | NEDD9 | 0.14020623 |
| 13882.p1 | 17:29559852:C:G | NF1 | 0.17245022 |
| 14131.p1 | 7:26224578:CCTTT:C | NFE2L3 | 0.16203686 |
| 13670.p1 | 1:61553905:C:T | NFIA | 0.13958744 |
| 14385.p1 | 9:14150143:C:T | NFIB | 0.15568171 |
| 11808.p1 | 6:31525439:C:CTG | NFKBIL1 | 0.13461243 |
| 13183.p1 | 14:51223326:AAT:A | NIN | 0.12373499 |
| 12036.p1 | 20:25459809:T:A | NINL | 0.15532108 |
| 11483.p1 | 17:43174508:ACCCGGCTGGCTCC:A | NMT1 | 0.13326214 |
| 13344.p1 | 9:139396487:C:T | NOTCH1 | 0.1672941 |
| 13197.p1 | 4:149073677:G:T | NR3C2 | 0.14488068 |
| 12501.p1 | 2:50724605:A:T | NRXN1 | 0.16821551 |
| 13349.p1 | 12:106461269:G:A | NUAK1 | 0.17164859 |
| 13839.p1 | 1:145589354:C:CCA | NUDT17 | 0.10836639 |
| 14473.p1 | 11:114441774:AC:A | NXPE4 | 0.12814231 |
| 14463.p1 | 1:102271635:GTTGT:G | OLFM3 | 0.25323209 |
| 12289.p1 | 1:158576487:G:GT | OR10Z1 | 0.14891236 |
| 11616.p1 | 1:248756275:G:T | OR2T10 | 0.14788699 |
| 12692.p1 | 11:4566664:A:T | OR52M1 | 0.14788699 |
| 11731.p1 | 12:76793728:TTCTTC:T | OSBPL8 | 0.1490981 |
| 14021.p1 | 17:3591305:C:T | P2RX5-TAX1BP3 | 0.13740831 |
| 12438.p1 | 11:117034592:C:G | PAFAH1B2 | 0.13153815 |
| 13982.p1 | 2:206480234:C:G | PARD3B | 0.18493674 |
| 12858.p1 | 9:37015070:AG:A | PAX5 | 0.13509709 |
| 12773.p1 | 5:140263886:CTTCG:C | PCDHA13 | 0.10295097 |

| | | | |
|---|---|---|---|
| 13018.p1 | 7:100201679:CT:C | PCOLCE | 0.13276853 |
| 13691.p1 | 20:17434474:AG:A | PCSK2 | 0.20248682 |
| 11707.p1 | 2:242795103:TG:T | PDCD1 | 0.14157058 |
| 11355.p1 | 2:239161576:G:A | PER2 | 0.14194097 |
| 14508.p1 | 1:207237157:AT:A | PFKFB2 | 0.15260597 |
| 13359.p1 | 21:45742909:C:CT | PFKL | 0.41733469 |
| 12323.p1 | 9:96439929:AT:A | PHF2 | 0.15543263 |
| 13903.p1 | 9:96437988:C:T | PHF2 | 0.1587783 |
| 11339.p1 | 11:45971023:CT:C | PHF21A | 0.37936593 |
| 14133.p1 | 6:64413433:CCG:C | PHF3 | 0.17475203 |
| 12383.p1 | 3:52454424:GC:G | PHF7 | 0.10623543 |
| 13162.p1 | 1:20966445:C:T | PINK1 | 0.11512679 |
| 13199.p1 | 2:219501023:C:CCATG | PLCD4 | 0.32738065 |
| 14107.p1 | 1:150129151:C:CGT | PLEKHO1 | 0.22097497 |
| 14189.p1 | 9:140358800:G:A | PNPLA7 | 0.1411814 |
| 13398.p1 | 1:151377903:T:TCGTCATCA | POGZ | 0.20316922 |
| 13627.p1 | 1:151378489:G:A | POGZ | 0.20316922 |
| 13333.p1 | 19:630058:G:A | POLRMT | 0.15000948 |
| 12340.p1 | 17:58740376:G:A | PPM1D | 0.15233107 |
| 14550.p1 | 6:42975179:CT:C | PPP2R5D | 0.1523483 |
| 11164.p1 | 1:228033225:G:A | PRSS38 | 0.17930427 |
| 11842.p1 | 8:18730164:TC:T | PSD3 | 0.10936444 |
| 11365.p1 | 17:65343511:G:A | PSMD12 | 0.17205799 |
| 11256.p1 | 6:43109471:C:T | PTK7 | 0.2667671 |
| 11453.p1 | 4:87696594:G:A | PTPN13 | 0.20004341 |
| 11077.p1 | 8:61484622:C:T | RAB2A | 0.29228819 |
| 13125.p1 | 20:1210539:C:T | RAD21L1 | 0.10190045 |
| 12703.p1 | 17:17699836:AG:A | RAI1 | 0.16339777 |
| 13760.p1 | 2:109392193:TCAGA:T | RANBP2 | 0.2658718 |
| 11806.p1 | 13:50125566:ATC:A | RCBTB1 | 0.13027199 |

| | | | |
|---|---|---|---|
| 13972.p1 | 19:1818480:C:T | REXO1 | 0.15999688 |
| 13162.p1 | 6:72889391:T:TA | RIMS1 | 0.14315868 |
| 13497.p1 | 6:73102488:C:CA | RIMS1 | 0.23118344 |
| 14336.p1 | X:106145431:C:T | RIPPLY1 | 0.13777306 |
| 13688.p1 | 6:127607886:CAT:C | RNF146 | 0.17960808 |
| 13451.p1 | 17:78353516:T:C | RNF213 | 0.21142358 |
| 12056.p1 | 9:36375929:A:G | RNF38 | 0.15362322 |
| 11475.p1 | 8:10468066:AG:A | RP1L1 | 0.1601725 |
| 14301.p1 | 15:41828848:C:A | RPAP1 | 0.13607573 |
| 13222.p1 | X:20193403:G:C | RPS6KA3 | 0.1822339 |
| 14258.p1 | 17:41139288:C:T | RUNDC1 | 0.14098859 |
| 14667.p1 | 7:87339897:G:A | RUNDC3B | 0.16859801 |
| 14257.p1 | 15:75137487:GA:G | SCAMP2 | 0.139951 |
| 11114.p1 | 2:166210819:G:T | SCN2A | 0.14348761 |
| 11892.p1 | 2:166201379:C:A | SCN2A | 0.14707956 |
| 11563.p1 | 1:1222518:C:A | SCNN1D | 0.17631986 |
| 11712.p1 | 1:53416506:CTG:C | SCP2 | 0.1733279 |
| 13506.p1 | 11:9111272:C:A | SCUBE2 | 0.15465479 |
| 13602.p1 | 9:139304579:A:AT | SDCCAG3 | 0.10857538 |
| 12933.p1 | 18:42532022:C:CG | SETBP1 | 0.14901161 |
| 12565.p1 | 3:47098932:AT:A | SETD2 | 0.21110489 |
| 14581.p1 | 16:28877932:C:CG | SH2B1 | 0.1230077 |
| 12380.p1 | 11:70332092:CTG:C | SHANK2 | 0.16706676 |
| 14172.p1 | 11:116766981:A:AAAAT | SIK3 | 0.16584489 |
| 11925.p1 | 10:21805727:TG:T | SKIDA1 | 0.15193767 |
| 12939.p1 | 17:42399123:TCA:T | SLC25A39 | 0.12948913 |
| 13799.p1 | X:152958717:G:T | SLC6A8 | 0.25409224 |
| 12300.p1 | 14:23243580:G:A | SLC7A7 | 0.33650444 |
| 13190.p1 | 1:40875459:T:A | SMAP2 | 0.14499072 |
| 13147.p1 | 10:97098949:CCA:C | SORBS1 | 0.22669381 |

| | | | |
|---|---|---|---|
| 14165.p1 | 17:49072429:GATCTA:G | SPAG9 | 0.21056506 |
| 12611.p1 | 2:32339776:C:CA | SPAST | 0.16141556 |
| 14592.p1 | 11:64939990:C:T | SPDYC | 0.18779738 |
| 11160.p1 | 1:16255762:G:GA | SPEN | 0.14954061 |
| 12197.p1 | 5:147506589:TC:T | SPINK5 | 0.15007921 |
| 13882.p1 | 2:54857033:G:T | SPTBN1 | 0.17429742 |
| 13857.p1 | 16:30745033:TGACAGCGACTG:T | SRCAP | 0.20201429 |
| 14147.p1 | 10:17738824:CT:C | STAM | 0.17707224 |
| 14613.p1 | 11:72468902:G:A | STARD10 | 0.2163476 |
| 12617.p1 | 15:42967936:C:T | STARD9 | 0.11339041 |
| 13851.p1 | 3:67578618:G:A | SUCLG2 | 0.10161126 |
| 12763.p1 | 10:104263992:C:A | SUFU | 0.12055627 |
| 12864.p1 | 11:67938481:C:T | SUV420H1 | 0.22283015 |
| 13073.p1 | 6:33408650:CTT:C | SYNGAP1 | 0.17871054 |
| 14136.p1 | 17:61482569:C:T | TANC2 | 0.12240661 |
| 13010.p1 | 3:100015120:G:C | TBC1D23 | 0.16061674 |
| 14612.p1 | 3:176752047:TA:T | TBL1XR1 | 0.16048413 |
| 11480.p1 | 2:162273322:AC:A | TBR1 | 0.16105057 |
| 13796.p1 | 2:162275481:A:AC | TBR1 | 0.14657439 |
| 12090.p1 | 10:114910883:G:A | TCF7L2 | 0.18850378 |
| 13069.p1 | 10:114901076:G:A | TCF7L2 | 0.18792656 |
| 14540.p1 | 11:121060495:T:TGGAG | TECTA | 0.27127342 |
| 14620.p1 | 16:69404384:A:G | TERF2 | 0.14845687 |
| 13647.p1 | 7:108204755:TAG:T | THAP5 | 0.19439088 |
| 13042.p1 | 7:11487022:G:A | THSD7A | 0.13446215 |
| 14266.p1 | 17:60655892:CAT:C | TLK2 | 0.14789348 |
| 12840.p1 | 3:196053904:C:T | TM4SF19 | 0.10284758 |
| 14467.p1 | 1:32542858:C:T | TMEM39B | 0.19783578 |
| 12340.p1 | 1:12175787:G:A | TNFRSF8 | 0.14484393 |
| 12052.p1 | 7:5347862:G:A | TNRC18 | 0.25074879 |

| | | | |
|---|---|---|---|
| 12673.p1 | 22:40661586:AAGAG:A | TNRC6B | 0.13866122 |
| 14163.p1 | 22:40662713:C:T | TNRC6B | 0.13866122 |
| 14086.p1 | 1:179820223:GGAGGT:G | TOR1AIP2 | 0.15179824 |
| 13586.p1 | 1:228596934:T:TG | TRIM17 | 0.18787163 |
| 12867.p1 | 2:230701696:G:A | TRIP12 | 0.13262451 |
| 12826.p1 | 1:193045125:ATTAG:A | TROVE2 | 0.13794343 |
| 11782.p1 | 9:73151650:AGT:A | TRPM3 | 0.21100001 |
| 11510.p1 | 11:2428988:C:T | TRPM5 | 0.22787221 |
| 11291.p1 | 5:176078840:C:A | TSPAN17 | 0.2362621 |
| 11457.p1 | 11:866614:AC:A | TSPAN4 | 0.40491534 |
| 14620.p1 | 9:25677688:C:T | TUSC1 | 0.14788699 |
| 12843.p1 | 2:200797909:G:A | TYW5 | 0.25774957 |
| 14401.p1 | 1:154227676:T:TC | UBAP2L | 0.28843024 |
| 12950.p1 | 7:138968839:CCTCT:C | UBN2 | 0.15414315 |
| 12161.p1 | 2:170732427:CAA:C | UBR3 | 0.13203833 |
| 14012.p1 | 8:103287805:A:AT | UBR5 | 0.26922371 |
| 14644.p1 | 2:234676519:C:T | UGT1A5 | 0.13506015 |
| 14530.p1 | 14:94088799:AC:A | UNC79 | 0.18384713 |
| 12463.p1 | 2:210745771:C:T | UNC80 | 0.14653385 |
| 11609.p1 | X:118985815:G:GA | UPF3B | 0.13612803 |
| 13672.p1 | 21:33757868:G:A | URB1 | 0.1272078 |
| 12521.p1 | 12:62775296:CTGAG:C | USP15 | 0.23073345 |
| 13054.p1 | 19:57641996:GA:G | USP29 | 0.15393927 |
| 12260.p1 | 6:99893695:G:T | USP45 | 0.14695794 |
| 12087.p1 | 1:108303413:AG:A | VAV3 | 0.14615215 |
| 13646.p1 | 9:35060455:TAGAAC:T | VCP | 0.14232936 |
| 11290.p1 | X:65247955:C:T | VSIG4 | 0.16245636 |
| 14200.p1 | 10:28879673:CAA:C | WAC | 0.11157681 |
| 14204.p1 | 10:28899668:AATTC:A | WAC | 0.13699917 |
| 12044.p1 | 4:85715777:G:A | WDFY3 | 0.13155805 |

| | | | |
|---|---|---|---|
| 13094.p1 | 4:85719152:T:A | WDFY3 | 0.12170605 |
| 14596.p1 | 2:128481973:AT:A | WDR33 | 0.18312034 |
| 13033.p1 | 22:46372611:C:T | WNT7B | 0.11094198 |
| 11042.p1 | 1:228111953:GC:G | WNT9A | 0.14250264 |
| 11415.p1 | 8:10755512:G:A | XKR6 | 0.12888254 |
| 14402.p1 | 4:69195952:AT:A | YTHDC1 | 0.20717247 |
| 12600.p1 | 19:47584867:CA:C | ZC3H4 | 0.1644185 |
| 12600.p1 | 19:47584862:GCT:G | ZC3H4 | 0.1644185 |
| 14486.p1 | 16:72831723:G:GT | ZFHX3 | 0.15375261 |
| 13009.p1 | 5:178359410:C:T | ZFP2 | 0.16447551 |
| 13176.p1 | 14:68272014:GC:G | ZFYVE26 | 0.14042181 |
| 13513.p1 | 10:293337:G:A | ZMYND11 | 0.19078984 |
| 14311.p1 | 19:12691751:G:GCACTTCA | ZNF490 | 0.15867139 |
| 12070.p1 | 16:2051648:G:A | ZNF598 | 0.14759359 |
| 14491.p1 | 19:52519608:CGA:C | ZNF614 | 0.15755836 |
| 14175.p1 | 15:66838947:T:TG | ZWILCH | 0.17020247 |
