## Supplementary Methods for "Exons as units of phenotypic impact for truncating mutations in autism"

SSC sequencing and phenotype data

We used exome sequencing and ASD probands' phenotypic data available in the Simons Simplex Collection (SSC) [1]. *De novo* LGD mutations were obtained from Iossifov *et al.*[2]. As a source of phenotypic data, we used the Prepared Phenotype Dataset (v15) from the SFARI Base online data portal (sfari.org/resources/sfari-base). The SSC inclusion/exclusion criteria, phenotype profiling instruments, and sequencing protocols have been described in previous publications[1,3-6].

To identify the effects of mutations on gene transcripts, we analyzed variants in SSC using Ensembl's Variant Effect Predictor (VEP) tool, which provides a series of annotations for each mutation based on its likely effect on gene transcripts (grch37.ensembl.org/info/docs/tools/vep)[7]. For each mutation in SSC, we used VEP to identify likely gene-disrupting (LGD) mutations by searching for the following predicted consequences: nonsense (translational stop gained) variants, frameshift indels, splice acceptor variants, and splice donor variants.

SVIP sequencing and phenotype data

We used sequencing and phenotypic data available in the Simons Variation in Individuals Project (SVIP) [8]. *De novo* LGD mutations and phenotype scores were obtained from Simons VIP Phase 2 Single Gene Dataset v4.0, available from SFARI Base online data portal (sfari.org/resources/sfari-base).

Adjustment of phenotype scores for gender and age

SSC and SVIP include ASD probands of both genders spanning a broad range of ages and phenotypic abilities. Some phenotypes, such as IQ scores (available for SSC only) and the Vineland Adaptive Behaviour Scales (available for both SSC and SVIP), are already normalized to account for developmental differences, and hence were suitable for direct comparison. In contrast, other phenotypes are available in the dataset as raw scores and often vary substantially with the age and/or gender of the proband. To account for the effects of age and gender, we adjusted raw scores for unnormalized phenotypes. Following an approach previously used by Buja, *et al*.[9], we adjusted phenotypic scores using linear regression. Specifically, using phenotypic scores from all probands in SSC, we performed a multivariate linear regression for each raw score, with age and gender as independent regression parameters. We used a binary variable (taking value 0 for male or 1 for female) to capture mean differences across genders. We then used the regression residuals as adjusted scores for comparing proband phenotypes.

Notably, while the SSC dataset does not provide the exact age of probands at the time when each specific phenotype was collected, we approximated the age of the probands by using the age at which the Autism Diagnostic Observation Schedule was administered (database column "age at ADOS").

Comparison of phenotype variability

To compare phenotypes between probands, we computed the absolute difference in IQ scores between individuals. We paired probands based on whether or not they are affected by similar LGD mutations. In our analyses, we compared pairs of probands: (1) with LGD mutations in the same gene, (2) with mutations in the same gene and within 1000 bp, (3) with mutations in the same exon, and (4) with mutations in the same exon and of the same gender. For each group of paired comparisons, we estimate the phenotypic variability by calculating the average absolute difference in IQs observed across all pairs. To estimate the statistical significance of differences in phenotypic variability, we then used one-tailed Mann-Whitney $U$ tests to compare pairwise differences.

Chromosomal distance between mutations

We computed the distance in base pairs between a pair of mutations in the same gene by taking the absolute difference between their chromosomal positions. For deletion mutations that affect a genomic interval (i.e. deletion of multiple base pairs), we used the chromosomal coordinates that lead to the smallest distance between mutations.

For example, the distance between a deletion and an upstream mutation was calculated based on the 5' coordinate of the deleted sequence, while the distance to a downstream mutation was calculated based on the 3' coordinate. Our results remained essentially unchanged when alternate distance metrics were used (not shown).

Protein sequence analyses

To calculate the position of ASD mutations in protein sequence, we mapped genomic coordinates onto their respective protein-coding sequences. Gene annotation data, including both genomic coordinates and the corresponding coding sequence intervals, were obtained from the Ensembl database using the BioMart interface (biomart.org)[10]. We then calculated the distances between mutations based on peptide positions. We then calculated correlations between relative peptide positions of mutations and the corresponding proband phenotypes.

Due to alternative splicing, the peptide distance between mutations can vary across transcript isoforms. In the analyses presented in the paper, we analyzed the mean peptide distance across all isoforms. However, we also used other definitions of distance, including: the maximal distance, the distance in the longest transcript isoform of the gene (or "canonical" isoform), or the median distance. Our results were consistent across different measures of peptide distance.

Distance-matched permutation tests

We performed a test in which we controlled for the peptide-sequence proximity of mutations in the same exon. To produce random pairs of mutations with peptide distances similar to those observed for mutations in the same exon, we used a rejection sampling approach. The motivation for this approach is that we were able to easily sample pairs of mutations in the same gene (by randomly choosing observed pairs of mutations). However, we wanted the peptide distances between mutations to match a distance distribution similar to the distances between mutations in the same exon.

Thus, we calculated two log-normal peptide distance distributions: a target distribution $t(d)$ (mean: 1.89, SD: 0.55) describing the distribution of distances between mutations in the same exon, and a sampling distribution $s(d)$ (mean: 2.50, SD: 0.54) describing the distribution of distances between mutations in the same gene, but not necessarily the same exon. Then, for a randomly selected pair of mutations sampled with replacement, we accepted it as a null distribution sample with acceptance probability:

$$p(\text{accept}) = \frac{t(d)}{m \cdot s(d)} \qquad (1)$$

Where $m$ is a constant that bounds the likelihood ratio between the distributions. The resulting rejection sampling algorithm generates samples approximating the target distribution $t(d)$ using proposals from the sampling distribution $s(d)$.

We then estimated the statistical significance of our results by comparing the observed differences in IQ to those observed for distance-matched sets of mutations in the null distribution.

GTEx genotype and expression data

To investigate the dosage effects across human tissues resulting from protein-truncating variants, we used data from the Genotype-Tissue Expression (GTEx) project[11]. The data were obtained via NCBI's Database of Genotypes and Phenotypes (dbGaP) (ncbi.nlm.nih.gov/gap). Specifically, we downloaded genotype data, raw RNA-seq data, and processed expression data summarized to genes, isoforms, and exons (Study Accession: phs000424.v6.p1). For RNA-seq data, we obtained SRA submitted files; these files contained the binary alignment map (BAM) for each RNA-seq experiment. In the analysis, we considered samples from ten major human tissues: adipose (subcutaneous), tibial artery, brain, heart (left ventricle), lung, skeletal muscle, tibial nerve, skin (not sun-exposed), thyroid, and whole blood.

In the downloaded genotype data, we identified small indels (≤ 6 bp) and single-nucleotide variants that were called using exome sequencing of 180 individuals (v6 from June 2014). Sequencing and variant calling protocols were previously described in Melé *et al.*[12]. To explore the dosage effects resulting from heterozygous variants, we included only heterozygous genotypes in our analysis. To limit the number of false positive genotypes, we only used calls with quality ≥ 20 (Phred-scale) for SNVs, and ≥ 30 for indels. In addition, to minimize the potential for downstream mapping and short read alignment biases, we excluded SNVs with non-unique flanking regions, based on UCSC mappability tracks. For SNVs, we excluded variants with 50 bp mappability less than one. For indels, we excluded variants with 36 bp mappability less than one. We limited our study to variants with allele frequency ≤ 0.05.

Quantification of allele-specific expression (AE)

AE for GTEx variants was quantified using RNA-seq data. In general, we followed previously developed protocols based on re-alignment of reads to local sequences containing either wild-type (WT) or variant alleles (tllab.org/data-software)[13]. For each individual, we first identified heterozygous genotypes and then extracted the flanking (±100 bp) reference genome sequences around each variant to produce a set of local sequences containing WT alleles. Next, we substituted the variant allele into each sequence to reflect the genotyped truncating variant, producing a matching set of local sequences with variant alleles. Short reads from RNA-seq experiments from an individual were then realigned separately against the reference and alternate sequences. To quantify the expression of each allele, we counted the number of reads aligning uniquely and without error to the WT and variant sequences, respectively. These numbers were then used as allele counts reflecting AE.

To extract flanking reference sequences, we used the fastaFromBed tool from the bedtools software suite (v.2.23.0)[14]. All reference sequences were taken from the human reference genome (hg19) provided by GTEx (gtexportal.org). For aligning short reads, we used BWA (v.0.7.3)[15], and kept only alignment calls with base quality ≥ 10 and mapping quality ≥ 35. Following the methods in Rivas *et al.*[13], we restricted our analysis to samples with at least 8 mapped reads, and to variants with median allele frequency, across all tissues, ≤ 0.95 and ≥ 0.05.

Gene expression changes due to LGD variants in GTEx

To quantify altered gene expression due to an LGD variant, we considered the changes in expression ($\Delta x$) compared to wild type as a combined effect of allele-specific expression (AE), alternative splicing (AS), and nonsense-mediated decay (NMD). To account for AE, we reasoned that only a fraction of total mRNA would be transcribed from each allele. To account for alternative splicing, we reasoned that transcripts would be spliced into multiple transcript isoforms, only some of which would retain the exon with the truncating mutation. Finally, we assumed that nonsense-mediated decay is an imperfect degradation process, in which some fraction of LGD-containing mRNA escapes NMD. Formally, we represented a change in expression as:

$$\Delta x = f \cdot \epsilon \cdot x_{\mathrm{exon}} \qquad (2)$$

where the parameter $f$ (ranging from 0 to 1) quantifies the fraction of total transcription from the allele harboring the LGD variant, the parameter $\epsilon$ (ranging from 0 to 1) quantifies NMD efficiency, and $x_{\mathrm{exon}}$ represents the wild-type expression level of transcripts with the LGD-containing exon, i.e. transcripts susceptible to NMD.

Because only post-NMD expression levels are experimentally observed, the relationship between measured and wild-type expression levels can be expressed as:

$$x'_{\mathrm{exon}} = x_{\mathrm{exon}} - \Delta x \qquad (3)$$

where $x'_{\mathrm{exon}}$represents the experimentally observed expression. Combining equations (2) and (3), we can express the effects of NMD in terms of $x'_{\mathrm{exon}}$:

$$\Delta x = \frac{f \cdot \epsilon}{1 - f \cdot \epsilon} \cdot x'_{\mathrm{exon}} \qquad (4)$$

In order to estimate $\Delta x$ for each gene, we needed to infer the parameters $f$ and $\epsilon$, which quantify AE and NMD efficiency respectively. As we describe in the following sections, we inferred these parameters probabilistically by fitting appropriate distributions. Notably, because we were interested in comparing the effects of NMD across different tissues, and since the efficiency of NMD may vary across tissues, we performed separate analyses for each tissue.

Parametric inference of AE

To model the expected distribution of $f$ for each gene, we considered the AE observed for synonymous variants in the gene because such variants are unaffected by NMD. We considered a hierarchical model in which the distribution of $f$ can be modeled as a beta distribution ($f \sim \mathrm{Beta}(\alpha_f, \beta_f)$), with hyperparameters $\alpha_f$ and $\beta_f$. In the model, each specific measurement of a synonymous variant's AE represents a binomial sample with parameter $f$ drawn from the beta distribution: $k \sim \mathrm{Binomial}(n, f)$, where $n$ is the total number of reads from both copies of the gene, and $k$ is the total number of reads from the synonymous variant allele. In this formulation, the inference of the underlying parameters can be performed by fitting a standard hierarchical beta-binomial model[16,17]. To model the distribution of $f$ in a given gene, we used the maximum *a posteriori* estimators for the beta distribution:

$$\hat{\alpha}_f, \hat{\beta}_f = \underset{\alpha_f, \beta_f}{\mathrm{argmax}} \prod_{v \in V_{\mathrm{syn.}}} \prod_{i \in C_v} \left[ \int_0^1 P(k_{v,i} \mid n_{v,i}, f) \cdot P(f \mid \alpha_f, \beta_f) \cdot df \right] \cdot P(\alpha_f, \beta_f) \qquad (5)$$

Here, $v$ indexes a specific synonymous variant from the set of all such variants $V_{\mathrm{syn.}}$, and $i$ indexes a specific individual carrying the variant allele, where $C_v$ is the set of all carriers. $k_{v,i}$ and $n_{v,i}$ are, respectively, the number of reads mapped to the synonymous allele $v$ and the total number of reads mapped in an AE experiment.

To perform the maximization in equation (5), we used a standard re-parameterization of $(\alpha_f, \beta_f)$ into $u = \frac{\alpha_f}{\alpha_f + \beta_f}$ and $v = \frac{1}{\sqrt{\alpha_f + \beta_f}}$ [18]. The parameters $(u, v)$ represent the mean and approximate standard deviation of the Beta distribution. To find values that maximize the posterior probability, we divided the parameter space into a discrete grid and used a grid search algorithm. For the hyperprior distributions, we chose a uniform distribution for the mean and a commonly used half-Cauchy distribution for the variance[19].

Note that the resulting estimators are based on multiple synonymous variants, each of which can have multiple individual carriers in the study population. As a result, the variance of the fitted distributions accounts for differences across both haplotypes and individuals. To minimize undersampling effects, we limited our analysis to genes with at least four different synonymous mutations and at least ten individual carriers.

Parametric inference of NMD efficiency

To infer NMD efficiency ($\epsilon$), we compared the observed AE for LGD variants in GTEx to the expected AE estimated based on synonymous variants in the affected genes. The relative difference between the observed and expected values was then used to quantify the efficiency of NMD. Since experimental measurements for LGD variants are relatively sparse (one experiment per carrier of each LGD variant), we used an empirical Bayes approach to fit a hierarchical model. Specifically, we first used the observed AE for all LGD variants to infer the distribution of NMD efficiency across variants, and then we estimated the likely efficiency of NMD for specific variants.

We assumed that NMD would function with some efficiency $\epsilon \in [0,1]$, and modeled the distribution of $\epsilon$ using a Beta distribution ($\epsilon \sim \mathrm{Beta}(\alpha_e, \beta_e)$). In this model, each AE measurement for an LGD variant $v$ corresponds to a binomial sample ($k \sim \mathrm{Binomial}(n, p)$), where $k$ is the number of reads with the LGD variant, $n$ is the total number of reads, and $p(f, \epsilon)$ is the underlying sampling parameter which depends on the AE parameter $f$ of the LGD allele and the NMD efficiency $\epsilon$. $p(f, \epsilon)$ can be expressed as:

$$p(f, \epsilon) = \frac{f \cdot (1 - \epsilon)}{(1 - f) + f \cdot (1 - \epsilon)} = \frac{f \cdot (1 - \epsilon)}{1 - f \cdot \epsilon} \qquad (6)$$

Where the numerator expresses the amount of post-NMD mRNA containing the LGD allele, and the denominator expresses the total amount of post-NMD mRNA from both alleles.

To learn the distribution of $\epsilon$ across variants, we searched for parameter values that would maximize the posterior probability given all experimental observations. Formally, the maximization can be written:

$$\hat{\alpha}_e, \hat{\beta}_e = \underset{\alpha_e,\beta_e}{\operatorname{argmax}} \prod_{v \in V_{\mathrm{LGD}}} \left[ \iint_{f,\epsilon} P_v\big(\, k \mid n, p(f,\epsilon) \,\big) \cdot P_v\Big(\, f \mid \hat{\alpha}_f, \hat{\beta}_f \,\Big) \cdot P\big(\, \epsilon \mid \alpha_e, \beta_e \,\big) \cdot df \cdot d\epsilon \right] \cdot P(\alpha_e, \beta_e) \qquad (7)$$

Here, $v$ indexes a specific LGD variant drawn from the set of all analyzed LGD variants ($V_{\mathrm{LGD}}$). The term $P_v\big(\, k \mid n, p(f,\epsilon) \,\big)$ represents the likelihood to observe AE for an LGD variant $v$. However, since $p(f,\epsilon)$, the binomial rate parameter, depends on $\epsilon$ and $f$ (see Eqn. 6), calculation of each likelihood requires integration across both parameters. Overall, the procedure searches for a distribution for $\epsilon$ that best explains deviations of AE for LGD variants from the expected AE distribution for each affected gene. The distribution of $\epsilon$ gives the likely percent decrease in expression due to NMD.

To perform the optimization, we re-parameterized the beta distribution using mean and variance parameters $u = \frac{\alpha_e}{\alpha_e + \beta_e}$ and $v = \frac{1}{\sqrt{\alpha_e + \beta_e}}$. Then we divided the parameter space into a discrete grid and calculated the posterior probability at each value of $(u, v)$. For the hyperprior distributions, we chose a uniform distribution for the mean and a half-Cauchy distribution for the variance parameter. We calculated the double integrals in Equation (7) numerically using Newton-Cotes cubature.

Using the maximum *a posteriori* values for parameters $(\alpha_e, \beta_e)$, we then inferred estimates of NMD efficiency for individual LGD variants. For each variant, we searched for maximum *a posteriori* estimates of $\epsilon$:

$$\begin{aligned} \hat{\epsilon} &= \underset{\epsilon}{\operatorname{argmax}}\, P\big(\, k \mid n, p(\epsilon) \,\big) \cdot P\big(\, \epsilon \mid \hat{\alpha}_e, \hat{\beta}_e \,\big) && (8) \\ &= \underset{\epsilon}{\operatorname{argmax}} \left[ \int_0^1 P\big(\, k \mid n, p(f,\epsilon) \,\big) \cdot P\big(\, f \mid \hat{\alpha}_f, \hat{\beta}_f \,\big) \cdot df \right] \cdot P\big(\, \epsilon \mid \hat{\alpha}_e, \hat{\beta}_e \,\big) && (9) \end{aligned}$$

The prior term was calculated directly from the parameters $\hat{\alpha}_e$ and $\hat{\beta}_e$. To calculate the total likelihood of $(k, n)$, which depends on rate parameter $p$, we integrated over possible values of $f$. The maximization in Equation (9), performed for each LGD variant separately, estimates the efficiency of NMD for a specific variant.

In our implementation, we estimated values of $\hat{\epsilon}$ using a one-dimensional parameter-scanning algorithm. For each value of $\epsilon$, we evaluated the integral over $f$ numerically, using Simpson's rule quadrature.

<u>Changes in total gene expression due to LGD variants</u>

Statistical inference of the parameters $f$ and $\epsilon$, described in the previous sections, allowed us to compute the expected changes in expression due to each LGD variant (see Eqn. 2). To account for differences in absolute expression level across genes, we normalized the effects of NMD by the total expression of affected genes:

$$\Delta x_{\mathrm{rel}} = \frac{\Delta x}{x_{\mathrm{gene}}} = \frac{\Delta x}{x'_{\mathrm{gene}} + \Delta x} \qquad (10)$$

Where $x_{\mathrm{gene}}$ is the total amount of mRNA expressed and $x'_{\mathrm{gene}}$ is the experimentally measured mRNA expression. In this way, $\Delta x_{\mathrm{rel}}$ expresses the normalized change in target gene expression due to NMD. In the manuscript, we compared $\Delta x_{\mathrm{rel}}$ either between pairs of variants in the same exon or in the same gene but different exons.

<u>Isoform-specific expression changes due to LGD variants</u>

To quantify the effect of LGD variants on splicing isoforms of a gene, we calculated the change in expression for each transcriptional isoform separately. Specifically, for a gene with *n* isoforms, we calculated the change in expression as (see Eqn. 2):

$$\Delta x_{\text{isoform } k} = \begin{cases} f \cdot \epsilon \cdot x_{\text{isoform } k} & \text{if isoform } k \text{ contains affected exon} \\ 0 & \text{otherwise} \end{cases} \quad (11)$$

To explore the effect of LGD mutations across all protein-coding isoforms of a gene, we represented the overall profile of isoform expression changes as an *n*-dimensional vector:

$$\bar{z} = (\Delta x_{\text{isoform } 1}, \cdots, \Delta x_{\text{isoform } k}, \cdots, \Delta x_{\text{isoform } n}) \quad (12)$$

To quantify the difference between isoform-specific expression change vectors, we used an angular distance metric between expression change vectors $\bar{z}_1$ and $\bar{z}_2$:

$$d(\bar{z}_1, \bar{z}_2) = \frac{2}{\pi} \cdot \cos^{-1} \frac{\bar{z}_1 \cdot \bar{z}_2}{\|\bar{z}_1\|_2 \cdot \|\bar{z}_2\|_2} \quad (13)$$

The dot ( $\cdot$) operator in the numerator represents the dot product between vectors. The denominator is the scalar product of the $\ell^2$ (Euclidean) norms for each vector. In this way, the distance $d$, which takes values on the unit interval $[0,1]$, computes a normalized angle between the expression change vectors for LGD variants. As with total gene expression in the manuscript, we compared the distances *d* either between pairs of variants in the same exon or in the same gene but different exons.

BrainSpan expression data

We obtained the human brain RNA-Seq expression data from the BrainSpan Atlas of the Developing Human Brain (brainspan.org)[20]. We used the BrainSpan project's developmental transcriptome RNA-seq dataset, which includes tables summarizing the average expression level in RPKM of human genes and their exons. RPKM values were quantile normalized across samples.

Correlation between changes in gene dosage and phenotypic effects

Human genes likely differ in their contributions to cognitive phenotypes. Therefore, for each gene with multiple LGD mutations in SSC, we estimated the IQ phenotype's sensitivity to changes in gene dosage (i.e. phenotype dosage sensitivity or PDS). To that end, we used least-squares linear regressions, regressing the observed phenotypic effects ($y$), defined as the difference between the average neurotypical IQ (100) and the proband's IQ, against the relative expression of LGD targeted exons $\left(x_{\text{rel}} = \frac{x_{\text{exon}}}{x_{\text{gene}}}\right)$. In each regression, we assumed that normal (wild type) gene dosage corresponds to a neurotypical IQ (100), and therefore fixed the *y*-intercept at 0. The slope ($s$) of the fitted least-squares regression line provided an estimate of the phenotypic sensitivity to gene dosage (Methods Figure 1).

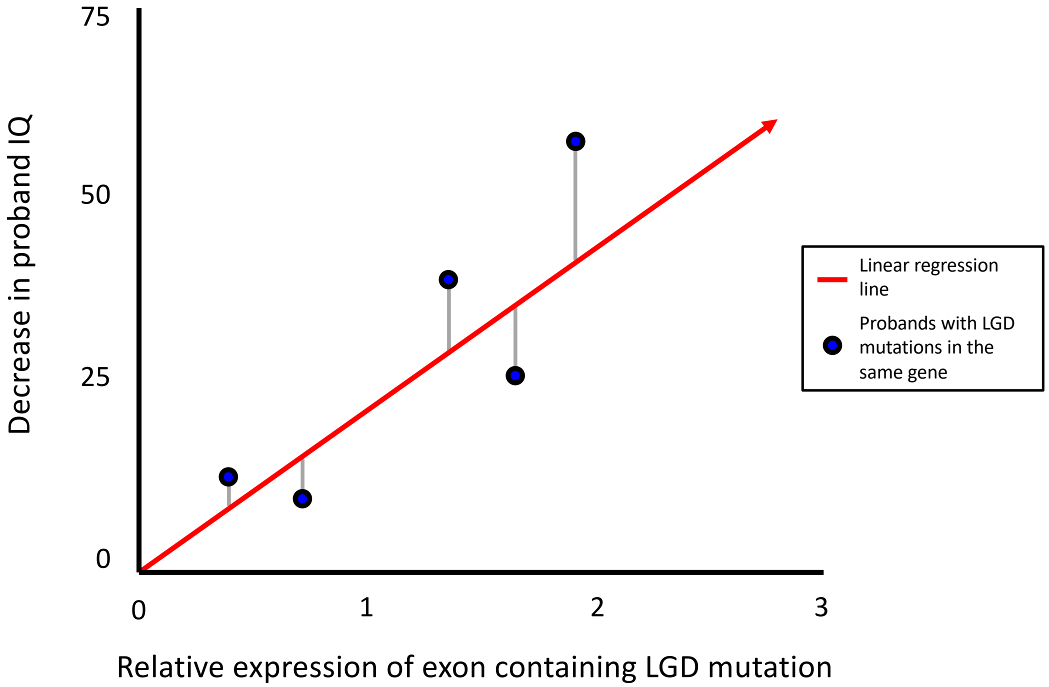


**Methods Figure 1:** An illustration of the method we used to estimate phenotypic sensitivity to changes in the dosage of a gene. For each gene with multiple truncating mutations in SSC, we used least-squares linear regression to estimate the phenotypic sensitivity. We defined phenotypic sensitivity as the slope of the fitted regression between the relative expression of the target exon $\left(\frac{x_{\mathrm{exon}}}{x_{\mathrm{gene}}}\right)$ and the effect of the mutation, i.e. the corresponding proband's IQ compared to the average neurotypical value (100). Each blue point in the figure represents a proband with an LGD mutation in the same gene. The *x*-axis position indicates the relative expression the targeted exon. The *y*-axis position indicates the observed effect of the mutation on IQ. The red line shows the least-squares regression line.

To express these linear model parameters in terms of gene dosage, we used experimental data from GTEx to model the relationship between the relative expression of an exon ($x_{\mathrm{rel}}$) and the corresponding dosage changes observed due to LGD mutation(s) in that exon. These predictions were made by averaging the dosage changes observed for GTEx LGD mutations in exons with similar relative expression, defined as values within the $\epsilon$ half-width interval $[x_{\mathrm{rel}} - \epsilon, x_{\mathrm{rel}} + \epsilon\,]$. We then averaged the observed dosage changes ($\Delta x_{\mathrm{obs}}$) for GTEx mutations in exons with relative expression values within the defined interval. Calculated in that way, predicted changes in dosage ($\Delta x_{\mathrm{pred}}$) as a function of relative expression can be expressed as:

$$\Delta x_{\mathrm{pred}}(x_{\mathrm{rel}}) = \frac{1}{|M|} \cdot \sum_{M} \Delta x_{\mathrm{obs}} \qquad (14)$$

Where $M$ is the set of LGD mutations in GTEx in exons with relative expression within $\epsilon$ of $x_{\mathrm{rel}}$. The parameter $\epsilon$ was set to 0.05, which was chosen as the interval for which the estimated averages for predicted dosage changes had SEM less than 1%.

We then analyzed phenotypic effects across multiple genes by normalizing $\left(y_{\mathrm{norm}} = \frac{y}{s}\right)$ the effect of each LGD mutation by the PDS ($s$) of the affected gene, defined as the expected decrease in IQ due to a 10% change in gene dosage. To establish the statistical significance of the correlation between normalized phenotypic effects and changes in gene dosage, we used a permutation test. Specifically, we reassigned LGD mutations to randomly selected exons in the same gene, with the reassignment probability proportional to the length of each exon. Then, following the same normalization procedure, we computed the correlation between normalized phenotypic effects and relative exon expression in the randomly shuffled data. The distribution of the correlations observed in the permuted data was used to estimate empirical *P*-values.

Linear model-based predictions of the effects of LGD mutations

Based on the dosage sensitivity model described in the previous section, we performed leave-one-out predictions of the effect ($y$) of each LGD mutation on full-scale, nonverbal, and verbal IQ, defined as the difference between the observed IQ and the average neurotypical IQ (100). To predict the effect of each withheld mutation, we performed least-squares linear regression using all other mutation in the same gene. In each regression, the phenotypic impact of mutations was the dependent variable, and the relative expression $\left(x_{\text{rel}} = \frac{x_{\text{exon}}}{x_{\text{gene}}}\right)$ of target exons was the independent variable. We assumed that mutations with no effect on gene dosage would result in neurotypical IQs and fixed the *y*-intercept of each regression at 0. The fitted regression lines (with slope $s$) were used to predict the phenotypic impact of the withheld mutation $\left(y_{\text{pred}} = s x_{\text{rel}}\right)$. Prediction errors were calculated as the absolute difference between predicted and observed effects $\left(\left|y_{\text{pred}} - y\right|\right)$.

Variance explained by same exon membership

To estimate the variance explained by same exon membership, we used LGD mutations in the same exon to infer IQ phenotypes for held out probands. Specifically, for each proband, we predicted that the phenotypic effect of an LGD mutation would be equal to the average effect for all other LGD mutations in the same exon. We then compared the variance explained by same exon membership to the variance explained by a multivariable linear model integrating various exon- and gene-level features, including prenatal and postnatal expression, exon evolutionary conservation, biological pathway membership, and protein domain truncation. We fitted a linear model using probands with LGD mutations in recurrently truncated genes, excluding probands with LGD mutations in the same exon as a testing set, then used the model to predict IQs for the held out probands. Same exon membership explained ~61% of IQ variance. The linear model integrating other features explained ~8% of variance.

Enrichment of mutations in developmentally biased exons

For each exon in BrainSpan, we calculated the developmental bias statistic, defined as the difference between mean prenatal and mean postnatal expression levels. Means were calculated for log-transformed values ($\log_2$ x+1) across all prenatal or postnatal samples. Only exons expressed in the brain were considered (average RPKM $\geq$1). Exons from genes harboring LGD mutations were partitioned into quartile groups based on developmental bias. Enrichments in each exon group were calculated by comparing the fraction of LGD mutations affecting the exons with fraction of coding sequence in the exons' coding sequences (CDS). Coding sequence lengths were obtained from Gencode v10 annotations.

Software availability

Most of the presented analyses, including hypothesis testing, data analysis, and statistical calculations, were performed using the R programming language (v.3.3.1 "Bug in Your Hair")[21], Python (v.3.5.2, Anaconda v.4.4.1), and Origin Pro 2017 (Version 94E; OriginLab, Northampton, MA). In addition, wherever other computational tools were used, we provided references to specific software packages used to perform the described analyses, including availability and versioning.

The quantification of AE, which we performed for specific synonymous and LGD variants in the GTEx dataset, required a specialized short-read analysis pipeline to perform reference sequence extraction, variant sequence construction, short read realignment, and AE quantification. These steps were implemented as a series of custom Python scripts; the scripts for this analysis are provided online.

We also wrote customized software to estimate the effects of nonsense-mediated decay. Specifically, we implemented numerical methods to fit distributions describing allele-specific expression and the efficiency of NMD. C source code for the software used to perform these calculations is also available online.

Data availability

Genotype and phenotype data from the SSC and VIP are available through the SFARI Base online portal (https://base.sfari.org/). Processed RNA-seq data from GTEx are available through an online data portal (https://www.gtexportal.org/); raw short read sequencing data are available through NCBI's dbGaP (accession number phs000424.v6.p1). The BrainSpan RNA-seq dataset is available online (http://www.brainspan.org/).

21 R Core Team. *R: A Language and Environment for Statistical Computing*. (R Foundation for Statistical Computing, 2013).
